# Pan-Cancer chromatin analysis of the human vtRNA genes uncovers their association with cancer biology

**DOI:** 10.1101/2020.10.05.324723

**Authors:** Rafael Sebastián Fort, María Ana Duhagon

## Abstract

The vault RNAs (vtRNAs) are a class of 84-141 nt eukaryotic non-coding RNAs transcribed by RNA polymerase III, named for their association with the conserved vault particle, a riboprotein complex whose function remains poorly understood. Of the 4 human vtRNA genes, the three clustered at locus 1, i.e. vtRNA1-1, vtRNA1-2 and vtRNA1-3, are integral components of the vault particle, while vtRNA2-1 is a more divergent homologue located in a second locus. Gene expression studies of vtRNAs in large cancer cohorts have been hindered by the failure of vtRNA sequencing using conventional transcriptomic approaches. However, since the vtRNAs transcription is regulated by DNA methylation, the analysis of the chromatin status of their promoters is a suitable surrogate approach to study their expression. Here we infer the landscape of vtRNA expression in cancer from the genome-wide DNA methylation (Illumina Infinium Human Methylation 450 K BeadChip) and chromatin accessibility (ATAC-seq) data of The Cancer Genome Atlas (TCGA). On average, vtRNA1-1 has the most accessible chromatin, followed by vtRNA1-2, vtRNA2-1 and vtRNA1-3. The correlation of the chromatin status of the vtRNA promoters and the binding sites of a common core of transcription factors stands for their transcriptional co-regulation by factors related to viral infection. Yet, vtRNA2-1 is the most independently regulated vtRNA homologue across tissue types. VtRNA1-1 and vtRNA1-3 chromatin status does not significantly change in cancer, though vtRNA1-3 promoter has repressive chromatin marks in a few cancer types. However, vtRNA2-1 and vtRNA1-2 expression are widely deregulated in neoplastic tissues and is compatible with a broad oncogenic role of vtRNA1-2, and both tumor suppressor and oncogenic functions of vtRNA2-1 depending of tissue contexts. Yet, vtRNA1-1, vtRNA1-2 and vtRNA2-1 promoter DNA methylation predicts a shorter patient overall survival cancer-wide. In addition, gene ontology analyses of co-regulated genes identifies a chromosome 5 regulatory domain controlling vtRNA1-1 and neighboring genes, and epithelial differentiation, immune and thyroid cancer gene sets for vtRNA1-2, vtRNA2-1 and vtRNA1-3 respectively. Furthermore, vtRNA expression patterns are associated with cancer immune subtypes. Finally, vtRNA1-2 expression is positively associated with cell proliferation and wound healing, in agreement with its oncogenic expression profile. Overall, our study presents the landscape of vtRNA expression cancer-wide, identifying co-regulated gene networks and ontological pathways associated with the different vtRNA genes that may account for their diverse roles in cancer.

## Introduction

The vault RNAs (vtRNAs) are a class of eukaryotic small non-coding RNAs (84-141 nt) transcribed by the RNA polymerase III, which associates to the vault particle (<5 % of the total mass of the vault particle) (Kedersha and Rome, 1986; Kedersha et al., 1991). Although this particle is the largest cellular nucleoprotein complex, its function remains scarcely understood. Most mammals have a single vtRNA gene (i.e., mouse and rat), but the human genome has three copies, annotated as vtRNA1-1 (98pb), vtRNA1-2 and vtRNA1-3 (88pb), which are located in the chromosome region 5q33.1 (the vtRNA1 locus) and transcribe the RNAs that are integral components of the vault particle. Another vault RNA gene in chromosome X, was classified as a pseudogene (vtRNA3-1P) due to the absence of expression in several cell lines and the presence of silencing mutations at the B box element of its promoter (van Zon et al., 2001). Lately, a transcript initially annotated as the human microRNA precursor hsa-mir-886 (Landgraf et al., 2007), was re-classified as another human vtRNA homologue and consequently renamed as vtRNA2-1 (Stadler et al., 2009). VtRNA2-1 is also located in chromosome 5 (5q31.1, locus vtRNA2) at 0.5Mb distance from the vtRNA1 cluster, where is placed between SMAD5 and TGFB1 genes in an antisense direction. Existing evidence indicates that vtRNA2-1 is neither associated with the vault particle nor co-regulated with the vtRNA1 locus (Lee et al., 2011; Stadler et al., 2009). Nevertheless, the realization that only 5-20% of all the cellular vtRNA transcripts are associated to the vault particle suggests additional roles for the majority of vtRNA transcripts, independent of the vault particle (Kickhoefer et al., 1998; Nandy et al., 2009; van Zon et al., 2001).

Most of the current knowledge of vtRNA function focuses on vtRNA1-1 and vtRNA2-1 roles in viral infection and cancer. Functional studies found that vtRNA1-1 and vtRNA1-2 (but not vtRNA1-3) can interact directly with drugs as mitoxantrone, doxorubicin and etoposide (Gopinath et al., 2005, 2010; Mashima et al., 2008). A recent report demonstrated that p62 protein interacts with vtRNA1-1 and inhibits the p62 dependent autophagy *in vitro* and *in vivo* (Horos et al., 2019). The vault RNAs have also been linked to native immune response, because of their strong upregulation during the infection of Influenza A virus (IAV) and Epstein-Barr virus (EBV) (Amort et al., 2015; Li et al., 2015; Nandy et al., 2009). These studies proposed a viral activation of host vtRNAs as a mechanism to maintain the inhibition of PKR signaling pathway, representing a viral strategy to circumvent host innate immunity.

Initial reports about vtRNA2-1 (locus vtRNA2) revealed that it is abundant in normal tissues while lowly expressed in cancer cell lines from different tissue origin (breast, melanoma, cervix, lung, oral and prostate), which is consistent with a tumor suppressive role in cancer (Lee et al., 2011; Treppendahl et al., 2012). Moreover, vtRNA2-1 was proposed as a new type of ncRNA functioning as a tumor suppressor gene (TSG) that inhibits PKR, and was consequently renamed as “nc886” (Golec et al., 2019; Jeon et al., 2012a, 2012b; Lee et al., 2011). Then, its anti-proliferative and TSG function was described in prostate (Aakula et al., 2015; Fort et al., 2018; Ma et al., 2020), skin (Lee et al., 2019a), gastric (Lee et al., 2014b) and esophageal (Im et al., 2020; Lee et al., 2014a) and cholangiocarcinoma cells (Kunkeaw et al., 2012). Conversely, a pro-proliferative and anti-apoptotic oncogenic role was proposed for vtRNA2-1 in renal (Lei et al., 2017), ovarian (Ahn et al., 2018), thyroid (Lee et al., 2016), cervical (Li et al., 2017) and endometrial (Hu et al., 2017) tissues. A recent finding proposing a vtRNA2-1/PKR loss mediated doxorubicin cytotoxicity introduced a novel view about its contribution to chemotherapy response (Kunkeaw et al., 2019). In addition, a potential TSG role has been put forward for vtRNA1-3 in Myelodysplastic syndrome (MDS) (Helbo et al., 2015).

Apart from their role as full length RNAs, vtRNAs are precursors of small RNAs. Indeed, small RNAs with microRNA like function were demonstrated to derive from vtRNA1-1 (svRNAs) and to repress the expression CYP3A4, a key enzyme of drug metabolism (Meng et al., 2016; Persson et al., 2009). VtRNA2-1 has also been shown to act as a microRNA precursor in different tissues, serving both as TSG in prostate (Aakula et al., 2015; Fendler et al., 2011; Fort et al., 2020), bladder (Nordentoft et al., 2012), breast (Tahiri et al., 2014), colon (Yu et al., 2011), lung (Bi et al., 2014; Cao et al., 2013; Gao et al., 2011; Shen et al., 2018) and thyroid cancers (Dettmer et al., 2014; Xiong et al., 2011) and as oncogene (OG) in renal (Yu et al., 2014), colorectal (Schou et al., 2014) and esophagus cancer (Okumura et al., 2016) through the repression of specific mRNA transcripts in human cancer.

The vtRNAs transcription is controlled by promoter DNA methylation. Different lines of evidence revealed that the epigenetic control of vtRNA2-1 is complex and owns clinical relevance in several tissues solid tumors including breast, lung, colon, bladder, prostate, esophagus, hepatic and stomach cancer (Cao et al., 2013; Fort et al., 2018; Lee et al., 2014a, 2014b; Romanelli et al., 2014; Treppendahl et al., 2012; Yu et al., 2020). Intriguing aspects of the epigenetic regulation of vtRNA2-1 locus comprise its dependence on the parental origin of the allele (Joo et al., 2018; Paliwal et al., 2013), and its sensitivity to the periconceptional environment ((Silver et al., 2015) and subsequent independent studies (van Dijk et al., 2018; Richmond et al., 2018)). In addition, vtRNA1-3 promoter methylation was associated with significantly poor outcome in lower risk myelodysplastic syndrome patients (Helbo et al., 2015).

Currently, transcriptomic sequencing is the benchmark technique to study global RNA expression (Stark et al., 2019). Nonetheless, some classes of RNAs are elusive to the standard transcriptomics due to their stable RNA structure and the presence of modified bases or ends, which impair the cDNA synthesis and/or adapter ligation during the sequencing library preparation (Sendler et al., 2011; Zheng et al., 2015). The vtRNAs belong to this group due to their conserved stable stem/hairpin loop secondary structure. The latter, together with the lack of sequencing of 40-200bp-long RNAs in the conventional transcriptomic studies, which are mostly intended for small and long RNAs, probably delayed their study in comparison with other regulatory RNAs. Nonetheless, since chromatin accessibility plays a critical role in the regulation of gene expression, at some extent the transcription of an RNA can be inferred from the chromatin status of its promoter (Corces et al., 2018; Duren et al., 2017; Liu et al., 2018). Since mounting evidence has shown that vtRNAs expression is tightly controlled by chromatin accessibility, dependent on nucleosome positioning and promoter DNA methylation (Ahn et al., 2018; Fort et al., 2018; Helbo et al., 2015, 2017; Lee et al., 2014a, 2014b; Park et al., 2017; Sallustio et al., 2016; Treppendahl et al., 2012), the chromatin structure is a suitable surrogate marker of vtRNA transcription and could be used as a proxy of their expression.

The expression, regulation, and role of the four human vtRNAs in normal and disease conditions are still poorly understood, and available knowledge indicates that they hold diverse tissue specific activities. The availability of genomic data of large sets of human tissues provided by The Cancer Genome Atlas (TCGA), allows the study of the human vtRNAs across 16 tissue types and normal/cancer conditions. Here, we analyze vtRNA genes chromatin of the TCGA patient cohort withdrawn from two approaches: the assay for transposase-accessible chromatin followed by NGS sequencing (ATAC-seq) to analyze chromatin accessibility (Buenrostro et al., 2013, 2015; Corces et al., 2018; Thurman et al., 2012) and the Illumina Infinium Human Methylation 450K BeadChip to analyze the CpG methylation of DNA (Moran et al., 2016). This led us to determine the patterns of transcriptional regulation of the four human vtRNAs throughout cancer tissues. We also evaluate the association between vtRNA expression and the patient clinical outcome in all the available cancer types. Finally, seeking for functional discoveries, we analyze vtRNA relations with immune subtypes and transcriptionally co-regulated gene programs. Our study reveals specific patterns of expression for each vtRNA that support previous evidence and poses potential new roles and molecular programs in which they may participate across and intra cancer types, increasing the comprehension of the role of the human vtRNAs in health and disease.

## Results

### VtRNA genes have different chromatin accessibility at the promoter region

The Cancer Genome Atlas consortium recently incorporated ATAC-seq data for tumor samples of the GDC Pan-Cancer dataset (385 primary tumors samples across 23 cancer types (Corces et al., 2018)). The ATAC-seq strategy is a genome-wide sequencing profiling approximation to chromatin accessibility using the hyperactive Tn5 transposase that simultaneously cut and ligate adapters preferentially to the accessible DNA regions. The counts of sequencing reads in a particular genomic region provides a direct measurement of chromatin accessibility (Buenrostro et al., 2015; Tsompana and Buck, 2014). For those gene promoters regulated by DNA methylation, ATAC-seq is a more direct approximation to chromatin accessibility than promoter CpG DNA methylation measurements because the latter is well correlated but not an exact measurement of chromatin accessibility. Indeed, the effect of DNA methylation on chromatin structure depends on the gene and the distribution of the methylation sites along it. It is also important to mention that gene expression interpretations based solely on chromatin accessibility are blind to post-transcriptional regulatory events such as processing, localization, and stability of the RNA transcript. Nevertheless, since the chromatin status of the different vtRNA promoters has been positively correlated with the abundance of their transcripts (Ahn et al., 2018; Fort et al., 2018; Helbo et al., 2015; Lee et al., 2014a, 2014b; Park et al., 2017; Sallustio et al., 2016; Treppendahl et al., 2012), ATAC-seq is likely a good surrogate of vtRNA expression. DNA CpG methylation data is available for 746 normal adjacent (across 23 tissues) and 8403 primary tumors (across 32 tissues) tissue samples of the Pan-Cancer TCGA cohort (Supplementary Table 1 and Supplementary Figure 1). The full data is listed in Supplementary Table 1 and the tissue composition of the three datasets of the cohort, comprising tumor ATAC-seq, normal and tumor DNA methylation, is shown in Supplementary Figure 1. The DNA methylation data has 9 more tissue types and a higher number of samples than the others, while the abundance of some tissues is skewed to twice enrichment in 1-4 specific dataset specific tissue types.

Although several reports demonstrated that the DNA methylation of vtRNA promoters is well correlated to vtRNA transcript expression in various tissues (Ahn et al., 2018; Fort et al., 2018; Helbo et al., 2015, 2017; Lee et al., 2014a, 2014b; Sallustio et al., 2016; Treppendahl et al., 2012), the association between the DNA methylation and chromatin accessibility at their promoters was not previously investigated.

To assess the expression of vtRNAs in cancer tissues, we analyzed a 500 bp region, containing the full vtRNA genes, the proximal promoter region and the RNA polymerase lII A and B box elements located downstream of the transcription start site (TSS) (Figure 1). Indeed, Vilalta et. al., 1994 demonstrated that the deletion of the 300bp 5’-flanking region of the rat vtRNA transcript largely reduced its transcription greater than 30-fold (Vilalta et al., 1994). As depicted in Figure 1, this region bears epigenetic marks of transcriptionally active chromatin (Li et al., 2007), including histone H3K27ac, H3K4me1, H3K4me3 at the promoters and DNaseI hypersensitivity clusters around the TSS in the cell types compiled by UCSC browser (Goldman et al., 2013). Interestingly, vtRNA2-1 is the only vtRNA immersed in a CpG island and vtRNA1-1 displays the strongest epigenetic marks of transcriptional activity (Figure 1). The analysis of the ATAC-seq values of this 500 bp region of all the TCGA primary tumors reveals that the four human vtRNAs have different levels of chromatin accessibility (Figure 2A). Moreover, the average DNA methylation of the promoters (Figure 2B) in primary tumor samples supports this finding (32 tissues, Supplementary Figure 1 and Supplementary Table 1). VtRNA1-1 promoter has the highest chromatin accessibility and the lowest DNA methylation, showing the smallest dispersion of values among the samples (Ave. 4.1, 0.6 SD) relative to the other three vtRNAs. Meanwhile, vtRNA1-2 (Ave. 2.9, 1.6 SD), vtRNA1-3 (Ave. 1.7, 1.2 SD) and vtRNA2-1 (Ave. 2.5, 1.4 SD) have broader ranges of ATAC-seq and DNA methylation values (Figure 2, Table 1, Supplementary Table 1 and Supplementary Figure 1). A comparison among the vtRNAs shows that the average chromatin accessibility of their promoters is inversely proportional to their average DNA methylation (Table 1 and Figure 2) (relative ATAC-seq values are vtRNA1-1 (4.1) > vtRNA1-2 (2.9) > vtRNA2-1 (2.5) and accordingly, relative DNA methylation are vtRNA1-1 (0.09) < vtRNA1-2 (0.4) < vtRNA2-1 (0.5)). Yet vtRNA1-3 average low chromatin accessibility is not accompanied by a comparatively denser promoter methylation (values 1.7 and 0.22 respectively). Nevertheless, the intra-sample correlation between the ATAC-seq and DNA methylation for the vtRNAs is between −0.24 and −0.78 (see rs-values in Table 1 and Supplementary Figure 2), suggesting that regardless of the relatively smaller average effect of DNA methylation on vtRNA1-3 promoter accessibility, all the vtRNAs are regulated by DNA methylation to some extent (Table 1). In addition, the CpG DNA methylation negatively correlates with ATAC-seq values in the 500bp of the promoters of the four vtRNAs (Table 1 and Supplementary Figure 2). The strength of the correlation for each vtRNA is higher for the two vtRNAs that have a broader spectrum of DNA methylation, i.e. vtRNA1-2 and vtRNA2-1 (rs = −0.74 and rs = −0.78 respectively) (Table 1 and Supplementary Figure 2B and 2D). Meanwhile, vtRNA1-1 and vtRNA1-3 promoters, which have narrower range of DNA methylation (vtRNA1-1 has also a small range of chromatin accessibility) present a smaller association between both values (rs = −0.24 and rs = −0.54 respectively) (Table 1 and Supplementary Figure 2A and 2C). Technical differences in ATAC-seq and DNA methylation array approaches, as well as naturally occurring non-linear correlation between average DNA methylation and chromatin structure may account for these differences (Corces et al., 2018; Liu et al., 2018). The low ATAC-seq value of the vtRNA3-1P pseudogene (Ave. −1.2, 0.6 SD), represents a proof of concept of the analyses (Supplementary Figure 3 and Supplementary Table 1).

**Figure 1.**
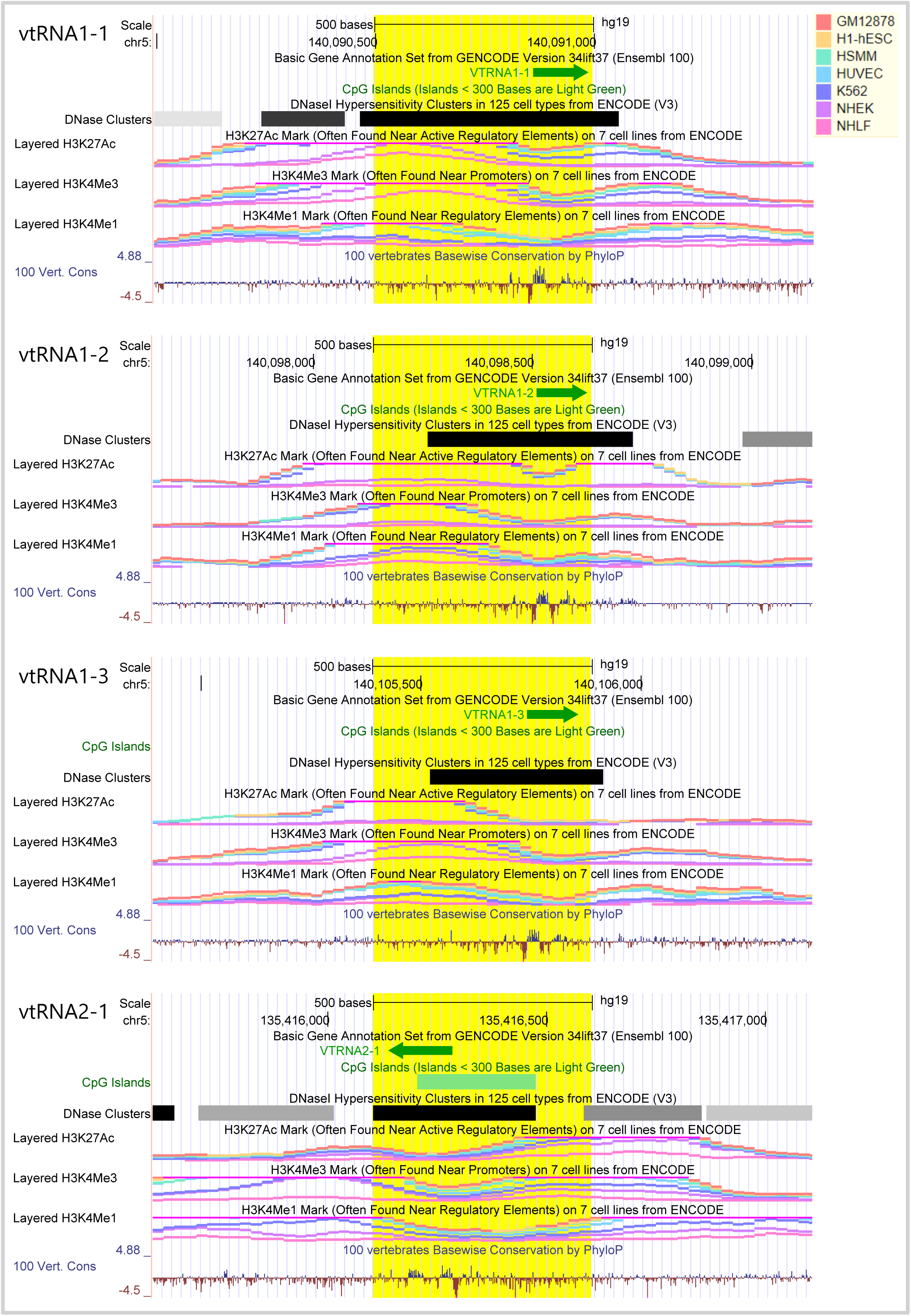
Genomic view of human vtRNA genes epigenetic features. Genomic view of 1.5 kb region of human vtRNA genes in UCSC Genome browser (GRCh37/hg19) centered at the 500bp bin highlighted in *yellow*, which was used for ATAC-seq and CpG methylation analyses. VtRNAs of locus 1 (vtRNA1-1, vtRNA1-2, vtRNA1-3) and of locus 2 (vtRNA2-1) clusters are in the sense and antisense orientation respectively. Several Gene annotation and ENCODE Project tracks for seven cell lines (GM12878, H1-hESC, HSMM, HUVEC, K562, NHEK, NHL) are displayed: DNA accessibility (DNaseI hypersensitivity clusters (color intensity is proportional to the maximum signal strength)), DNA methylation (CpG islands length greater than 200 bp), histone modification (H3K27Ac, H3K4me1, H3K4me3 marks), conservation of the region in 100 Vertebrates (log-odds Phylop scores). The vertical viewing range of the tracks displays the default settings of the browser for each variable.

**Figure 2.**
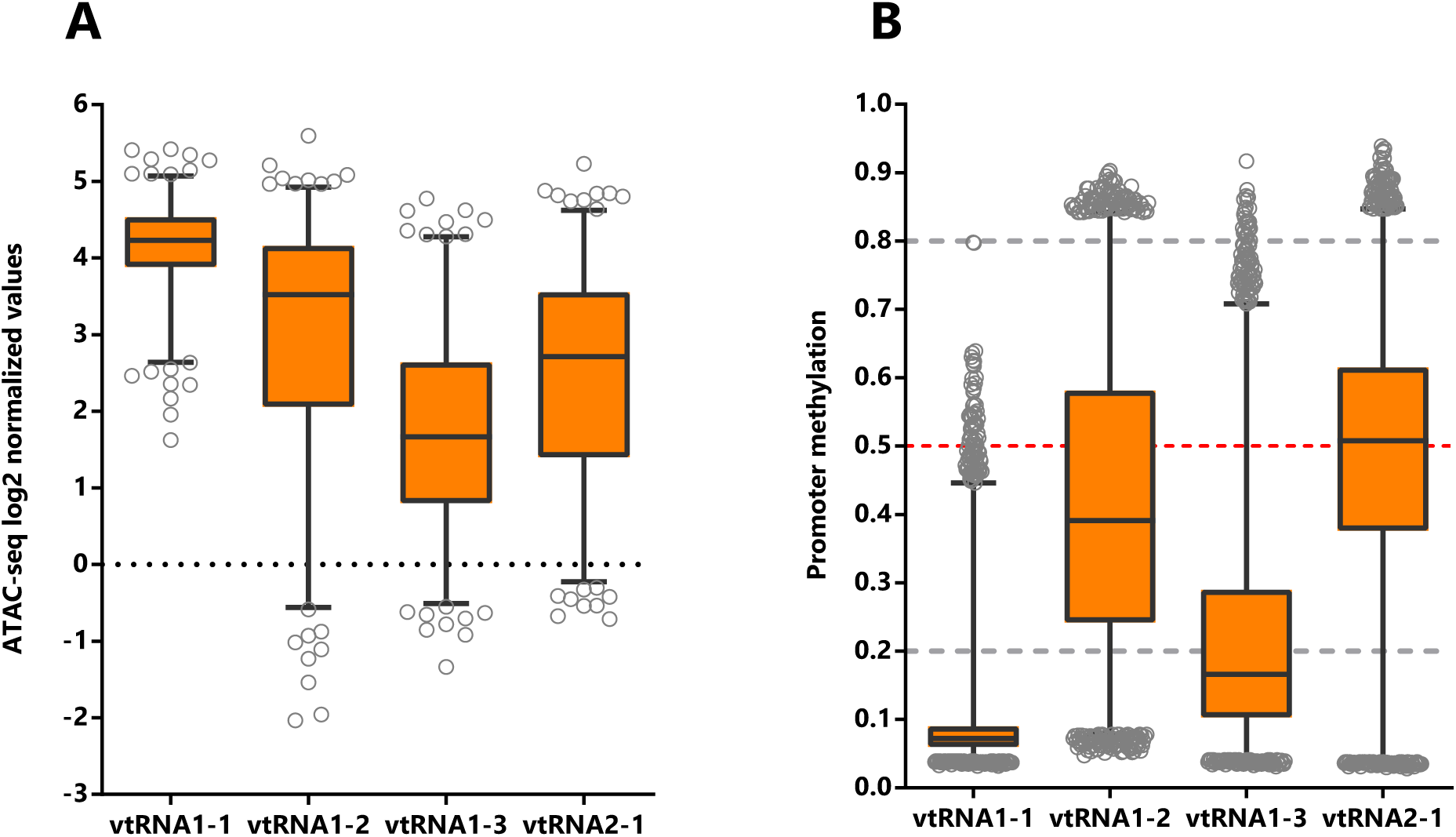
Chromatin accessibility and DNA methylation of the vtRNA promoters. Analysis of vtRNA 500pb promoter region of **A.** Average chromatin accessibility ATAC-seq values for the 385 primary tumors available at the Pan-Cancer TCGA dataset (23 tissues types) and **B.** Average DNA methylation beta-values for the 8403 primary tumors (32 tissues types). Dashed horizontal lines denote unmethylated (bottom gray, average beta-value ≤ 0.2), 50% methylated (middle red, average beta-value = 0.5) and highly methylated (top gray, average beta-value ≥ 0.8) promoter. The box plots show the median and the lower and upper quartile, and the whiskers the 2.5 and 97.5 percentile of the distribution.

**Table 1.**
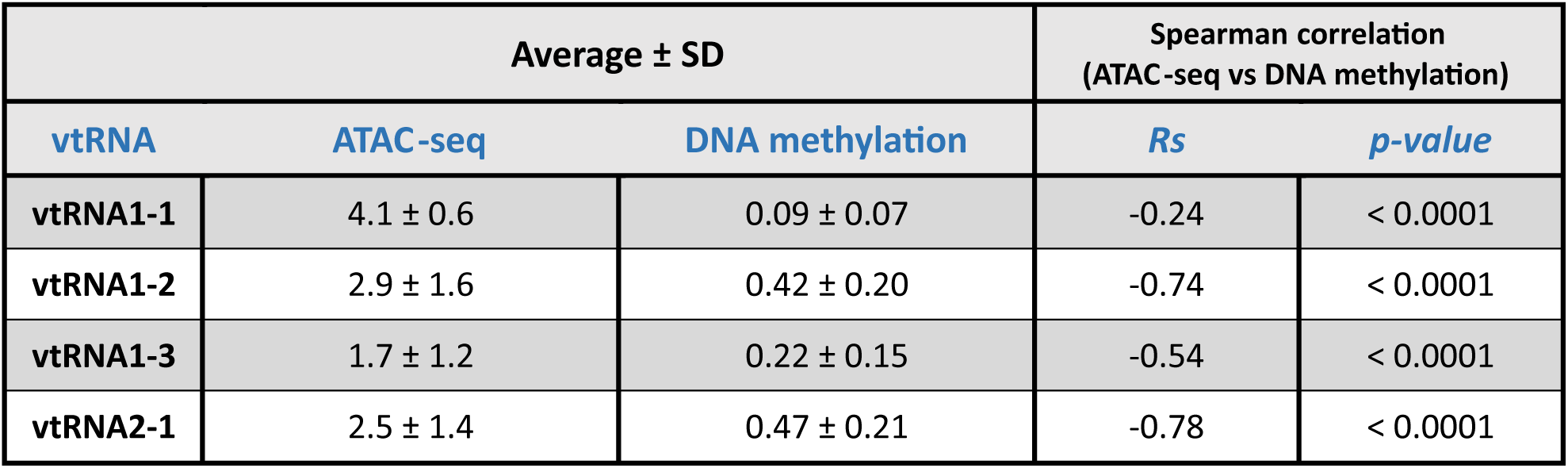
Correlation between vtRNA promoters ATAC-seq and DNA methylation in the Pan-Cancer TCGA dataset. The Spearman correlation was calculated for the 329 primary tumors studied by ATAC-seq and DNA methylation.

| Average ± SD |  |  | Spearman correlation<br>(ATAC-seq vs DNA methylation) |  |
| --- | --- | --- | --- | --- |
| vtRNA | ATAC-seq | DNA methylation | *Rs* | *p-value* |
| vtRNA1-1 | 4.1 ± 0.6 | 0.09 ± 0.07 | -0.24 | < 0.0001 |
| vtRNA1-2 | 2.9 ± 1.6 | 0.42 ± 0.20 | -0.74 | < 0.0001 |
| vtRNA1-3 | 1.7 ± 1.2 | 0.22 ± 0.15 | -0.54 | < 0.0001 |
| vtRNA2-1 | 2.5 ± 1.4 | 0.47 ± 0.21 | -0.78 | < 0.0001 |

### Chromatin accessibility, DNA methylation and transcription factor binding at the four vtRNA promoters are correlated

Aiming to investigate a possible co-regulation of vtRNA transcription, we determined the pairwise correlations in chromatin accessibility and DNA methylation between the four vtRNA promoters. We found that all pairs of vtRNAs, except vtRNA1-2 and vtRNA2-1, shown positive correlations in both datasets (Figure 3A and 3B) (rs-values 0.23-0.41 for ATAC-seq and rs-values 0.20-0.50 for DNA methylation). The unsupervised clustering of these epigenetic marks indicates that the three vtRNAs clustered at vtRNA1 locus (vtRNA1-1, vtRNA1-2 and vtRNA1-3), which are the vault particle associated vtRNAs, are more similar than vtRNA2-1 (Figure 3A and 3B).

**Figure 3.**
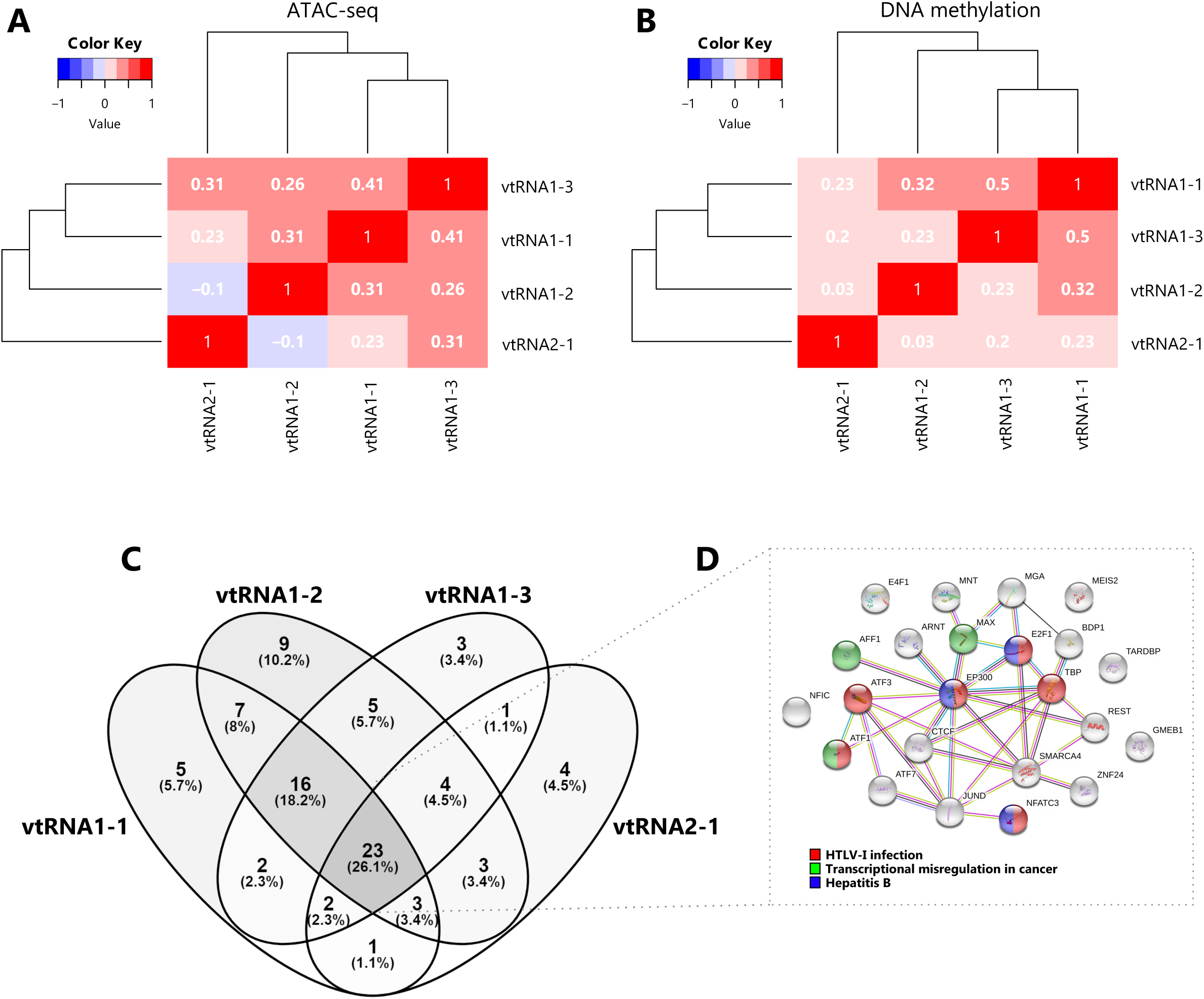
Comparative chromatin accessibility and transcription factor occupancy at the vtRNA loci. **A-B** Matrix of pairwise Spearman correlations (two-way hierarchical clustering distance measured by Euclidean and Ward clustering algorithms) of vtRNA 500pb promoter region for ATAC-seq data (385 primary tumors samples across 23 tissues) (A) and DNA methylation average beta-values data (8403 primary tumors samples across 32 tissues) (B). **C.** Venn diagram of transcription factors identified as ChIP-seq Peaks by ENCODE 3 project in the cell line K562 (Venny 2.1; https://bioinfogp.cnb.csic.es/tools/venny/index.html). The region for TFs assignment was defined as ±3000 pb from the vtRNA transcript sequence boundaries. **D.** Interaction cluster of the core 23 TFs common to all vtRNAs performed with STRING (Szklarczyk et al., 2017). The colored labels of the top 3 enriched KEGG pathway terms (FDR < 0.05) are indicated.

Given that the chromatin accessibility of a DNA region is partially controlled by transcription and chromatin remodeler’s factors (Corces et al., 2018; Klemm et al., 2019), we analyzed transcription factors (TFs) occupancy at a ± 3000 pb region centered at each vtRNA gene. Since there is no such study of the TCGA samples, we investigated the CHIP-seq data of the K562 chronic myelogenous leukemia cell line, which has the richest TFs data at ENCODE dataset (Goldman et al., 2013). We observe that the 4 vtRNAs share a common core of 23 TFs (26%). The 3 members of the vtRNA1 cluster share 16 additional TFs (18%), reaching 39 common TFs (44%), while only 2-4 TFs with vtRNA2-1 (Figure 3C and 3D and Supplementary Table 2). We finally asked whether this common core of TFs is linked to a specific biological process. The STRING enrichment analysis of the core of 23 TFs common to all vtRNAs identified HTLV-I infection (ATF1, ATF3, E2F1, EP300, NFATC3, TBP), Hepatitis B (E2F1, EP300, NFATC3) and Transcriptional misregulation in cancer (AFF1, ATF1, MAX) as the top 3 enriched KEGG pathway terms (FDR < 0.05) (Figure 3C and Supplementary Table 2).

Taken together these findings indicate that the transcriptional activity of the vtRNAs is gene specific, being vtRNA1-1 the most accessible and possibly the most expressed. In addition, the data suggest that the regulatory status of the vtRNA promoters is coordinately controlled, and the three vtRNA1s are more co-regulated among themselves. Yet, the core of TFs shared by the vtRNAs is associated with viral infection and cancer related terms in the myeloid cell line K562.

### The vtRNA promoters present gene and tissue specific patterns of chromatin accessibility in primary tumors

In order to investigate the expression of vtRNAs in different tissues we analyzed the Pan-Cancer TCGA ATAC-seq data discriminating the tissue of origin. As expected, vtRNA1-1 has the highest promoter chromatin accessibility among tissues and the smallest variation (Figure 4A, Supplementary Table 1 and Supplementary Figure 1). Remarkably, low-grade glioma tumor samples (LGG) show a global reduction in chromatin accessibility at the 4 vtRNA promoters (Figure 4A). An opposite pattern is seen in adrenocortical carcinoma tissue (ACC), where the vtRNAs have the highest concerted chromatin accessibility (Figure 4A). Individually, vtRNAs promoter accessibility is maximum and minimum in THCA (4.7) and LGG (3.3) for vtRNA1-1, ACC (4.2) and LGG (0.29) for vtRNA1-2, ACC (3.7) and LGG (0.24) for vtRNA1-3 and KIRC (4.2) and LGG (1.0) for vtRNA2-1 (Figure 4A). VtRNA3-1P reaches a maximum of promoter chromatin accessibility in Testicular Germ Cell Tumors (TGCT) (0.65) with an extremely low average ATAC-seq value and its minimum in PRAD (−1.5) (Supplementary Figure 3C). Remarkably, although vtRNA1-1 and vtRNA1-3 have the highest and lowest promoter accessibility in the majority of tissues respectively, vtRNA1-2 and vtRNA2-1 are the most variable (vtRNA1-1 SD = 0.34, vtRNA1-2 SD = 1.4, vtRNA1-3 SD = 0.75 and vtRNA2-1 SD = 0.78) (Figure 4A). In addition, in 12 tissues the chromatin accessibility of vtRNA1-2 is higher than vtRNA2-1 (57%) whereas in 9 tissues the chromatin accessibility of vtRNA1-2 is lower than vtRNA2-1 (43%) (Figure 4A).

**Figure 4.**
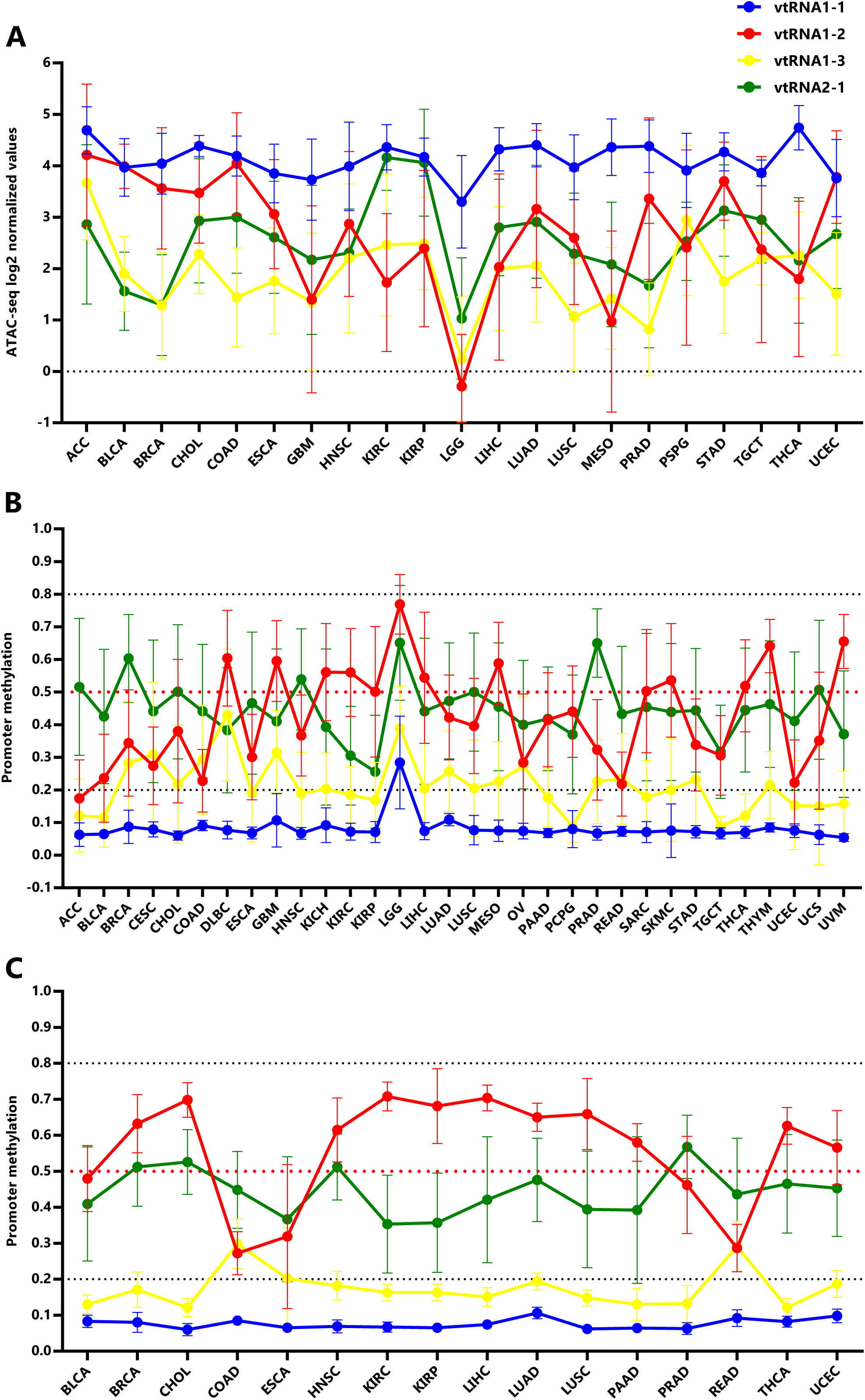
Chromatin accessibility and DNA methylation of the vtRNA promoters in tumor and normal tissues. **A.** ATAC-seq values for vtRNA promoters in 21 primary tumor tissues. **B.** Average beta-values of promoter DNA methylation for vtRNAs in 32 primary tumors tissues. **C.** Average beta-values of promoter DNA methylation for vtRNAs in 16 normal adjacent tissues. For **A**, **B** and **C**, the charts show the average value and standard deviation of each vtRNAs for the tissues with at least five samples available at Pan-Cancer TCGA dataset.

Using the vtRNAs promoter DNA methylation data, we extended the analysis to 32 primary tumor tissues, incorporating nine more tissues than ATAC-seq samples (DLBC, KICH, OV, PAAD, READ, SARC, THYM, UCS, UVM) (Figure 4B, Supplementary Figure 1) and 16 normal adjacent tissues (Figure 4C, Supplementary Figure 1). Again, the DNA methylation profile mirrors the chromatin accessibility in the individual tissue types (Figure 4). Remarkably, the association between the average vtRNA promoter chromatin accessibility and DNA methylation of the vtRNA1 cluster in each tissue type is higher than that observed for the average of all tissues (vtRNA1-1 rs = −0.28, vtRNA1-2 rs = −0.90 vtRNA1-3 rs = −0.74 and vtRNA2-1 rs = −0.58) (Table 1 and Supplementary Figure 4). The opposite finding for vtRNA2-1 may be explained by the impact of the previously recognized chromatin polymorphisms in the locus (Silver et al., 2015), which may prevail over the tissue specific variation for this gene.

In primary tumor tissues, promoter DNA methylation of vtRNA2-1 is higher than vtRNA1-2 in 17 tissues (53%) and the opposite is observed in 15 tissues (47%) (Figure 4B), while in normal tissues, promoter DNA methylation of vtRNA2-1 is higher than vtRNA1-2 in 4 tissues (25%) and the opposite is observed in 12 tissues (75%) (Figure 4C), indicating that although vtRNA1-2 is more methylated in normal samples, vtRNA2-1 gains methylation in neoplastic tissue, becoming more methylated than vtRNA1-2 (in 7 tissue types: BLCA, BRCA, CHOL, HNCSC, LUAD, LUSC, UCEC). Indeed, the Fisher’s exact test identifies a mirrored profile of chromatin accessibility between normal (4 tissues with high vtRNA1-2 and 12 tissues with high vtRNA2-1 from 16 total tissues) and tumor tissues (17 tissues with high vtRNA1-2 and 15 tissues with high vtRNA2-1 from 32 total tissues) for vtRNA1-2 and vtRNA2-1 (p-value = 0.07). Despite this deregulation in transformed tissues, we wondered if the tissue specific differences in vtRNA promoters were explained by their pre-existing status in the normal tissues. The correlation among average vtRNA promoter DNA methylation in normal and tumor tissues suggest that the variation in chromatin accessibility among tissue types is already established in the normal tissue counterparts (vtRNA1-1 rs = 0.53, vtRNA1-2 rs = 0.84, vtRNA1-3 rs = 0.47 and vtRNA2-1 rs = 0.75, Supplementary Figure 5).

### VtRNA promoter’s DNA methylation is associated with tumor stage and tissue of origin

The assessment of differential vtRNA expression from normal to tumor condition at the TCGA cohort can only be performed using DNA methylation dataset since no ATAC-seq analyses were performed in the normal tissues. The average beta-value of CpG sites in the 500bp vtRNAs promoter of normal and tumor tissue evidenced gene specific patterns of deregulation in neoplastic tissues (Figure 5). As depicted in Figure 4, vtRNA1-1 and vtRNA1-3 promoter DNA methylation and chromatin accessibility is low and high tissue-wide respectively, in both normal and tumor tissues (yet, there are highly methylated tumor outliers as LGG, that lack normal counterpart) (Figure 4 and Figures 5A and 5C). Conversely, vtRNA1-2 and vtRNA2-1 revealed significant differences in average promoter DNA methylation among the tissue categories (Figures 5B and 5D). In agreement with previous results the methylation of vtRNA1-2 promoter decreases from normal to tumor samples, while vtRNA2-1’s increases (Figures 5B and 5D and Figure 4).

**Figure 5.**
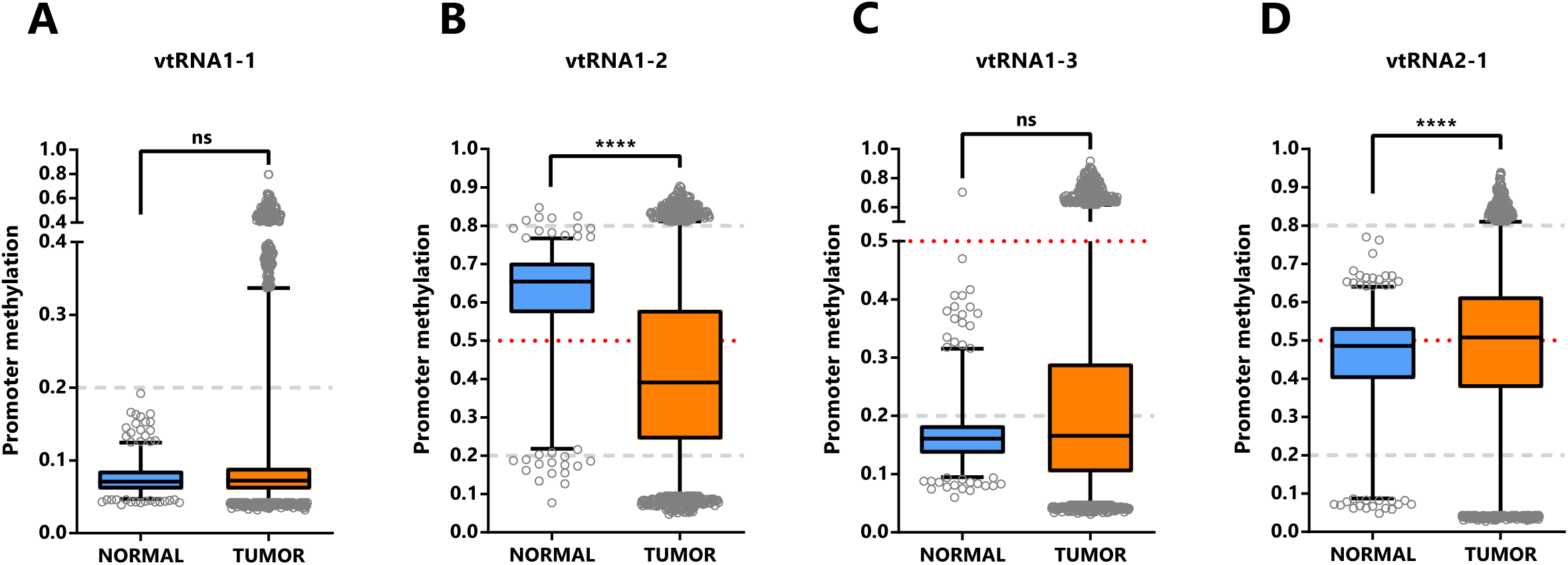
Promoter DNA methylation of vtRNAs in Normal and Tumor tissues of Pan-Cancer TCGA dataset. Average beta-values of promoter DNA methylation of vtRNA1-1 (**A**), vtRNA1-2 (**B**), vtRNA1-3 (**C)** and vtRNA2-1 (D), assessed in 746 normal and 8403 tumor tissues respectively. The box plots show the median line and lower and upper quartile and the whiskers the 2.5 and 97.5 percentile. Horizontal grey striped and red dotted lines denote unmethylated (average beta-value ≤ 0.2), 50% methylated (average beta-value = 0.5) and highly methylated (average beta-value ≥ 0.8) promoters. One-way ANOVA multiple test analysis with Sidak as posthoc was performed. **** p-value < 0.0001.

We next asked whether the promoter status of the vtRNAs is deregulated during the normal and tumor transformation of different tissue types. This comparison was restricted to the 16 tissues with normal samples (with data of at least five samples, Supplementary Figure 1, Supplementary Table 1 and Figure 6). The results revealed that vtRNA1-1 is globally unmethylated in normal and tumor from different tissue origins, but presents small but statistically significant changes in 4 tissue types (OG profile in BLCA, THCA UCEC and TSG in LUSC), whose biological impact remains to be determined (Figure 6A). Similarly, vtRNA1-3 is globally unmethylated, showing larger variability and significant upregulation in BRCA and PRAD, compatible with a TSG function (TSG trends for LIHC p-value = 0.08 and LUSC p-value = 0.09, Figure 6C). In agreement with the literature, vtRNA2-1 presents a TSG pattern of deregulation in PRAD, LUSC and BRCA (Cao et al., 2013; Fort et al., 2018; Romanelli et al., 2014), and an OG pattern in KIRP that has not been previously described (Figure 6D). Finally, a statistically significant dysregulation of vtRNA1-2 promoter DNA methylation across almost all cancer types (13 out of 16 tissues analyzed) poses it as a candidate OG in cancer (Figure 6B).

**Figure 6.**
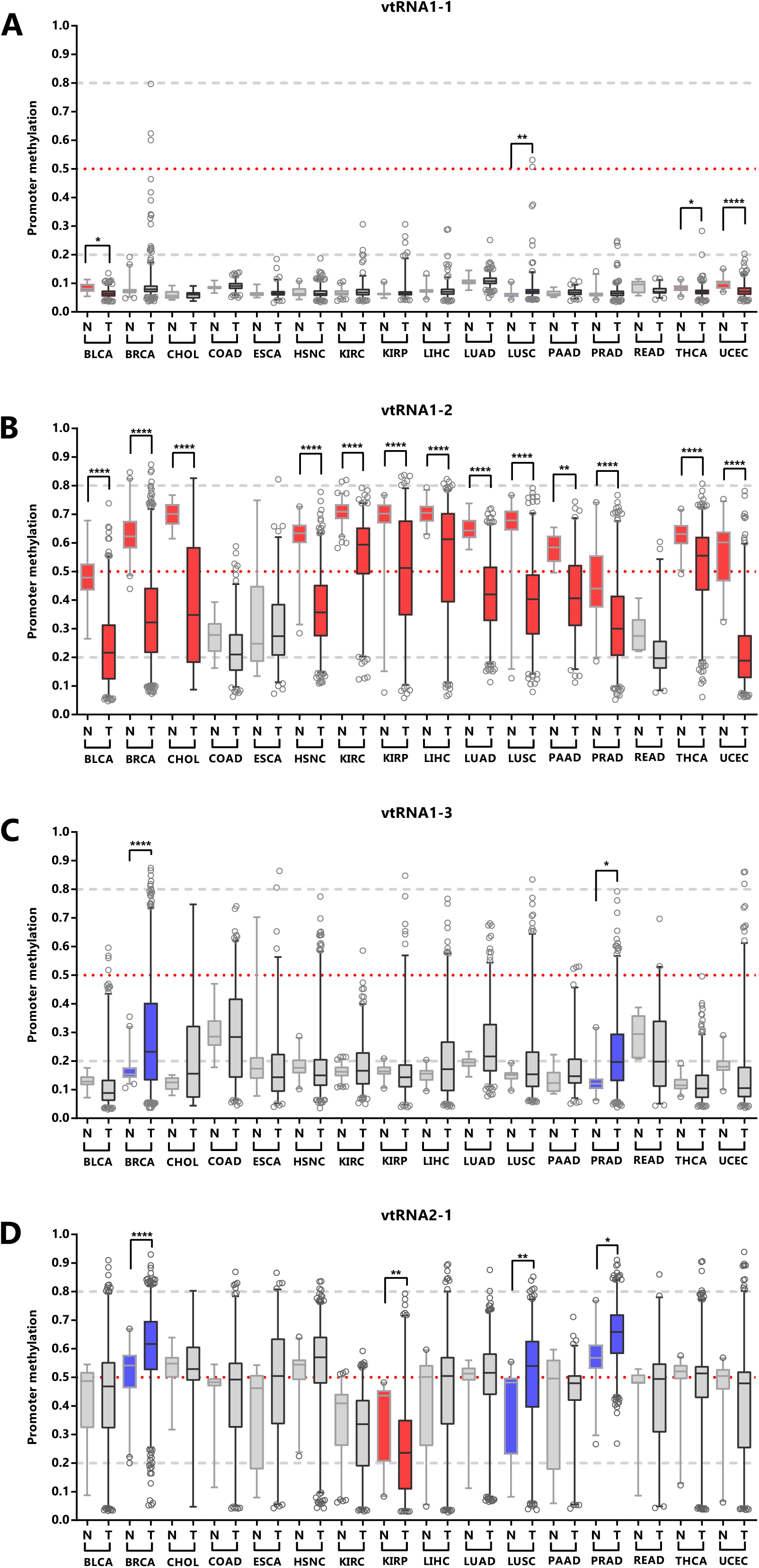
VtRNA promoters DNA methylation in Normal vs. Tumor samples. Average beta-values of promoter DNA methylation for vtRNA1-1 (**A**) vtRNA1-2 (**B**), vtRNA1-3 (**C**) and vtRNA2-1 (**D**). Acronyms indicate the tissue condition (normal (N) and tumor (T)). *Blue* and *red colors* indicate an increase or reduction in promoter methylation in tumor vs their normal tissues counterparts, compatible with a TSG and OG function respectively. The box plots show the median and the lower and upper quartile, and the whiskers the 2.5 and 97.5 percentile of the distribution. Horizontal, lines denote the methylation level of the promoters: grey striped bottom and top for unmethylated (average beta-value ≤ 0.2) or highly methylated (average beta-value ≥ 0.8) respectively, and red dotted for 50% methylated (average beta-value = 0.5)) promoters. One-way ANOVA multiple test analysis with Sidak as posthoc was performed. * p-value < 0.05; ** p-value < 0.01; **** p-value < 0.0001.

Overall, the data shows different DNA methylation changes in the vtRNAs promoters upon malignant transformation in several tissues. In addition, the deregulation of each vtRNA occurs mostly in a gene specific direction, where vtRNA1-2 has an OG like epigenetic de-repression while vtRNA2-1 (and to a lesser extent vtRNA1-1, vtRNA1-3) has a TSG like repressive methylation in tumor tissues.

### High chromatin accessibility at the promoter of vtRNA1-1, vtRNA1-3 and vtRNA2-1 is associated with low patient overall survival

Seeking for a possible clinical significance of the chromatin accessibility changes of the vtRNA promoters, we studied its association with the overall survival of patients (Figure 7 and Supplementary Table 3). The analysis of patients stratified by vtRNA promoter accessibility quartiles did not show differences in overall survival (data available at Supplementary Table 3). However, a significantly lower patient survival probability when patient tumors have lower promoter DNA methylation is observed for vtRNA1-1 (p-value = 0.003), vtRNA1-2 (p-value = 0.004) and vtRNA2-1 (p-value = 0.002) (patient stratified in quartiles of the promoter DNA methylation cohort values, Figure 7). Therefore, a lower promoter DNA methylation of vtRNA1-1, vtRNA1-2 and vtRNA2-1, a surrogate of their high expression, might be associated with poor patient overall survival cancer-wide.

**Figure 7.**
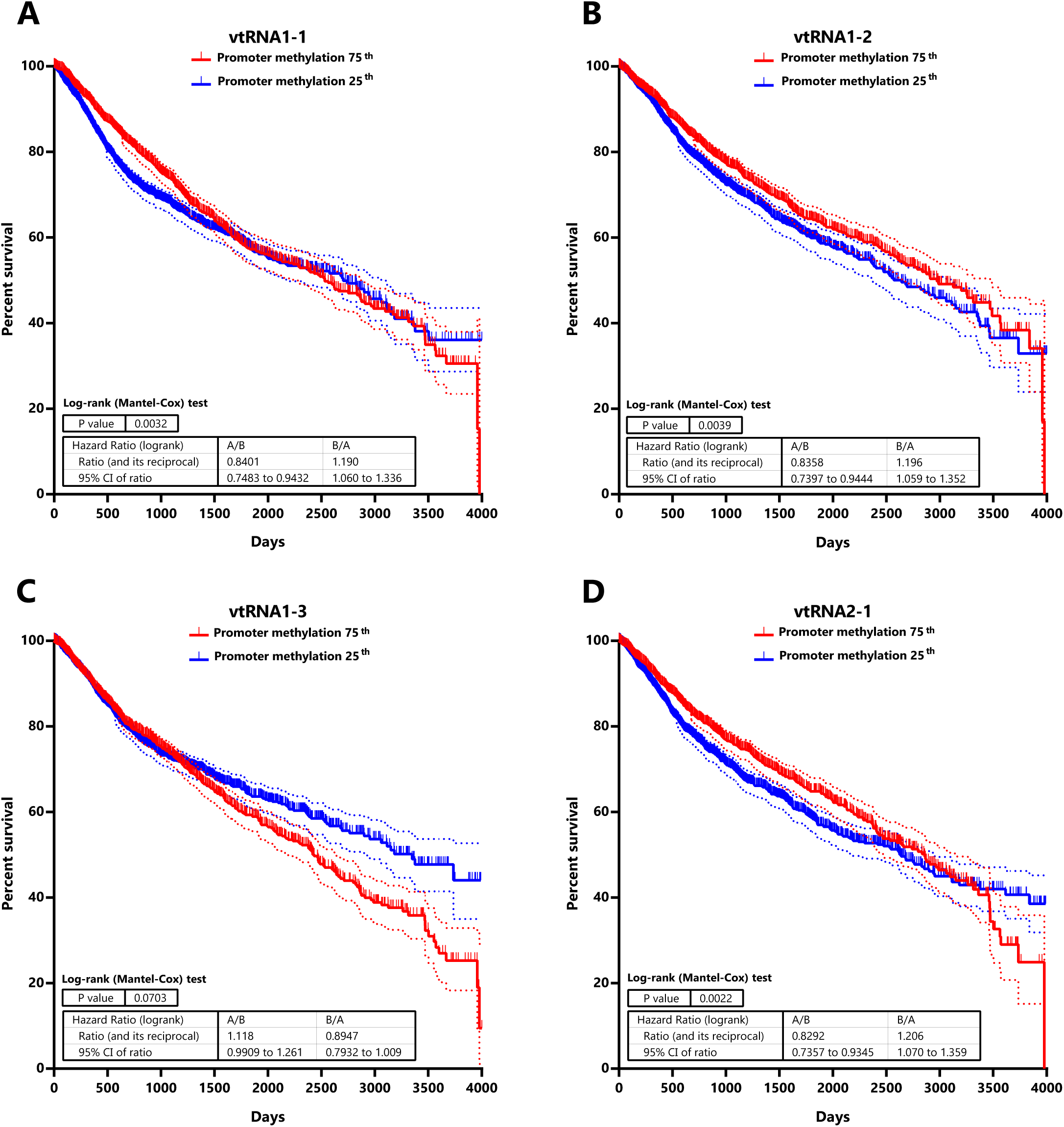
Patient Overall survival relative to vtRNA promoters DNA methylation status in Pan-Cancer TCGA dataset. Kaplan-Meier curves of overall patient survival probability over the time (4000 days) discriminating the primary tumors into two cohorts with relatively higher (75^th^) and lower (25^th^) DNA methylation at each vtRNA promoter. Promoter DNA methylation data of the 8060 primary tumors was used to stratify the patients in two quartiles, based on the expression of vtRNA1-1 (**A**) vtRNA1-2 (**B**), vtRNA1-3 (**C**) and vtRNA2-1 (**D**). Patient survival probability of the two groups was analyzed using Log-rank (Mantel-Cox) test. The dotted lines represent the 95% confidence interval for each curve.

### Genome-wide chromatin accessibility correlations with vtRNA promoters reveals their link to specific cancer related functions

Seeking to get insight into the function of the vtRNAs from their transcription regulatory marks, we searched for the genes co-regulated at chromatin level. We calculated the correlation of the ATAC-seq promoter values of each vtRNA and all the individual genes in the 385 primary tumor cohort and selected those showing a Spearman correlation rs ≥ 0.4. Using STRING software analysis, we then investigated their connection with Biological Process categories and KEGG pathways (Szklarczyk et al., 2017). A few enriched terms for the genes correlated with vtRNA1-2, vtRNA1-3 and vtRNA2-1 is identified (FDR < 1×10^-3^) (Supplementary Table 4). The 303 genes correlated with vtRNA1-2 are enriched in the Biological Process “Epithelial cell differentiation” (FDR < 1×10^-4^) (Supplementary Table 4). Meanwhile, the 550 genes correlated with vtRNA2-1 are enriched in the Biological Processes “Immune system process”, “Inflammatory response” and “Immune response” (FDR < 1×10^-5^) and the KEGG pathways term “Cytokine-cytokine receptor interaction” (FDR < 1×10^-4^) (Supplementary Table 4). The only 11 genes correlated with vtRNA1-3 are enriched in the “Thyroid cancer” KEGG pathway term (FDR < 1×10^-4^). Remarkably, although STRING analysis of the 46 vtRNA1-1 associated genes did not find enriched process or pathways, 45 of the 46 correlated genes are located at chromosome 5q region. Using the Cluster Locator algorithm (Pazos Obregón et al., 2018), we found a statistically significant non-random clustering behavior for the chromosomal location of the genes co-regulated with vtRNA1-1 (p-value < 1×10^-10^). Indeed, when the analysis was extended to 213 enes with Spearman correlation rs ≥ 0.3, 186 genes were situated at chromosome 5 (p-value < 1×10^-10^). These genes are positioned in particular regions at chr5q31, chr5q35, chr5q23, chr5q14, chr5q32, chr5q34 (FDR < 1×10^-4^) (Kuleshov et al., 2016) (Supplementary Table 4). We did not find a similar phenomenon for the other 3 vtRNAs. In summary, vtRNA1-3, vtRNA1-2 and vtRNA2-1 transcription may be co-regulated with genes belonging to specific biological processes located elsewhere, while vtRNA1-1 transcription may be controlled by a locally specialized chromatin domain at chromosome 5q region lacking apparent functional relatedness.

### Immune Subtypes associated with the vtRNAs profile

Taking into account that vtRNAs have been previously related to the immune response (Golec et al., 2019; Li et al., 2015; Nandy et al., 2009), and that we found an association of vtRNA2-1 with immune related terms, we sought to investigate a putative relation between the vtRNAs expression and the six Immune Subtypes defined by Thorsson et al. (Thorsson et al., 2018), compiled at Pan-Cancer TCGA (Supplementary Table 5). These six Immune Subtypes are named because of the foremost immune characteristic, comprising wound healing, IFN-g dominant, lymphocyte depleted, inflammatory, immunologically quiet and TGF-b dominant types (Thorsson et al., 2018). As was expected, mirrored patterns were observed for promoter ATAC-seq and DNA methylation data of the six groups (Figure 8). The immunologically quiet subtype shows a concerted shutdown of all vtRNA promoters (Figure 8), which is largely composed by LGG samples showing low vtRNA ATAC-seq and high promoter methylation average values in LGG tumors (Figures 4A and 4B). The immunological quiet subtype exhibits the lowest leukocyte fraction (very low Th17 and low Th2) and the highest macrophage:lymphocyte ratio (high M2 macrophages) (Thorsson et al., 2018). The remaining subtypes present a similar vtRNA profile except for the inversion of vtRNA1-2/vtRNA2-1 accessibility ratio, which is higher than 1 for the wound healing and IFN-g dominant subtypes while is lower than 1 for the lymphocyte depleted and inflammatory subtypes (Figure 8A). As expected, a mirrored pattern was observed for vtRNA1-2 and vtRNA2-1 at DNA methylation data (Figure 8B). Since the wound healing and IFN-g dominant subtype have high proliferation rate opposite to the lymphocyte depleted and inflammatory subtypes (Thorsson et al., 2018), we evaluated the correlation between vtRNA1-2 and vtRNA2-1 chromatin status and proliferation rate or wound healing status defined by Thorsson et al. (Supplementary Table 5) (Thorsson et al., 2018). A negative correlation between vtRNA1-2 promoter DNA methylation and the proliferation rate (rs = −0.44) and wound healing features of the tumors is revealed (rs = −0.48) (Supplementary Table 5). These findings suggest that vtRNA transcriptional regulation may be associated with tumor immunity.

**Figure 8.**
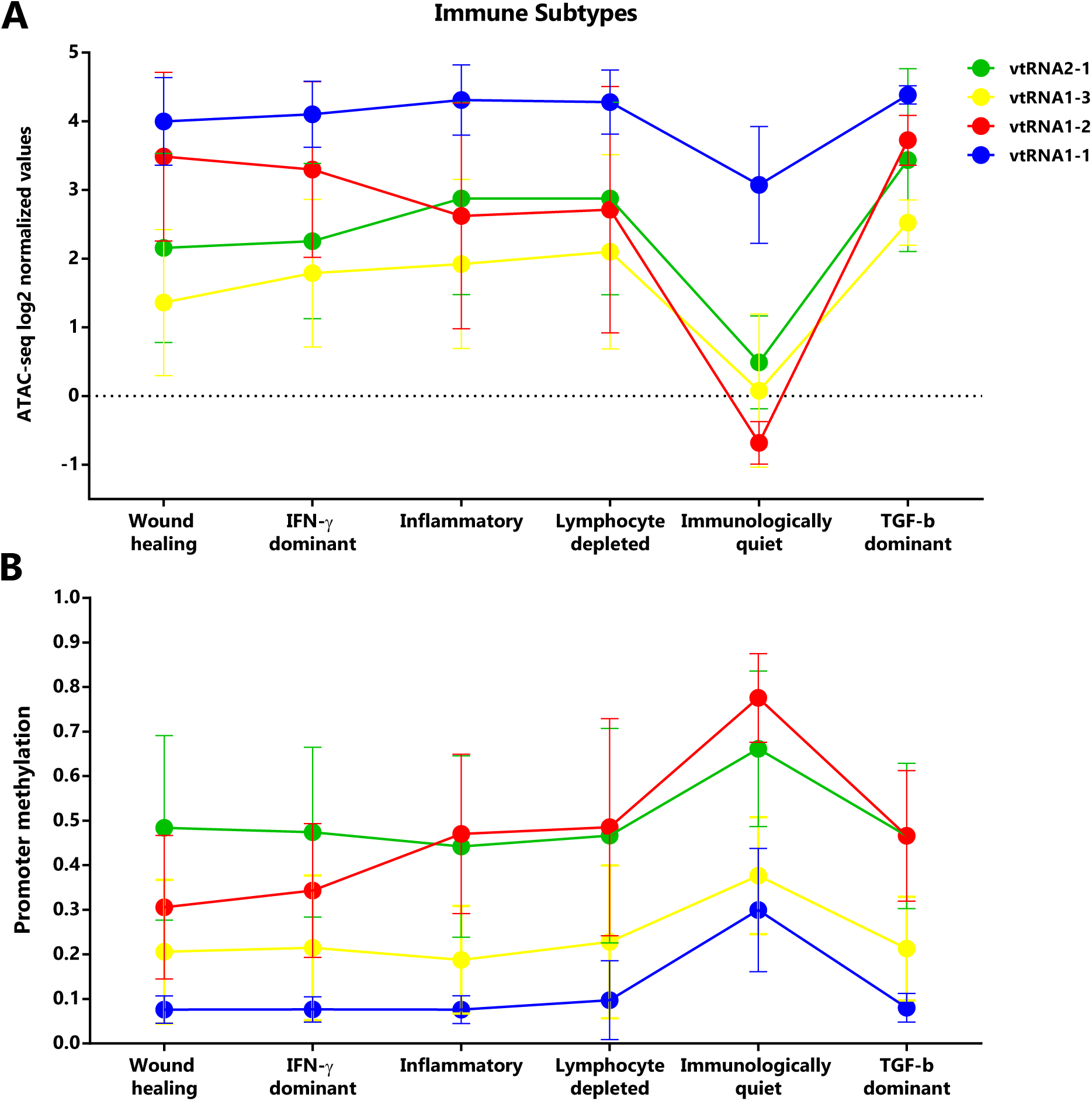
Chromatin accessibility and DNA methylation of vtRNA promoters in the Six Immune Subtypes analyzed in the PANCAN TCGA dataset. **A.** ATAC-seq values expressed as log2 normalized values for vtRNA promoters grouped in six Immune Subtype: wound healing (105 samples), IFN-g dominant (93 samples), inflammatory (110 samples), lymphocyte depleted (43 samples), immunologically quiet (9 samples), TGF-b dominant (4 samples). **B.** DNA methylation average beta-values for vtRNA promoters grouped in six Immune Subtype: wound healing (1963 samples), IFN-g dominant (2137 samples), inflammatory (2126 samples), lymphocyte depleted (933 samples), immunologically quiet (383 samples), TGF-b dominant (151 samples) (Thorsson et al., 2018). The charts show the average and standard deviation for each vtRNA promoter in each Immune Subtype group.

## Discussion

Although important insight into the vtRNA function has been recently gained, the landscape of vtRNA expression in human tissues and cancer was unknown, perhaps for the absence of vtRNA data in transcriptomic studies. Here we used ATAC-seq and methylation arrays data from the Pan-Cancer TCGA consortium (Weinstein et al., 2013), as surrogate variables to uncover the expression of vtRNAs across multiple cancer types. A limitation of our study is its blindness to post-transcriptional controls of vtRNA expression, such as RNA processing, transport, localization and/or stability. Indeed, there is evidence of vtRNA transcript regulation via RNA methylation by NSUN2, or via its association to other proteins like DUSP11, DIS3L2, SSB, SRSF2 and DICER (Burke et al., 2016; Hussain et al., 2013; Kickhoefer et al., 2002; Łabno et al., 2016; Persson et al., 2009; Sajini et al., 2019). Nonetheless, the chromatin status of the vtRNA promoters has been correlated with the expression of their transcripts in several reports (Ahn et al., 2018; Fort et al., 2018; Helbo et al., 2015, 2017; Lee et al., 2014a, 2014b; Sallustio et al., 2016; Treppendahl et al., 2012). These evidences unequivocally stand for the use of chromatin structure as a suitable proxy for vtRNA expression.

An integrated genomic view of the regulatory features of the vtRNA genes compiled in UCSC genome browser indicates that the 4 vtRNAs have chromatin signs of active transcription in different cell lines and defines a proximal promoter regulatory region of approximately 500pb (Nikitina et al., 2011; Vilalta et al., 1994). High resolution chromatin accessibility mapping studies by NOMe-Seq of the vtRNA promoters in cell lines validate this assumption (Helbo et al., 2017). Particularly, vtRNA2-1 is singular for being immersed in a CpG island, for lacking the TATA box, DSE or PSE elements characteristic of the others vtRNA promoters, and for bearing a cAMP-response (CRE) element (Canella et al., 2010; Helbo et al., 2017; Nandy et al., 2009). Since ATAC-seq approach is currently the best high throughput method to determine the chromatin accessibility of a DNA region directly (Chen et al., 2016; Sun et al., 2019; Tsompana and Buck, 2014), we interrogated the ATAC-seq data of TCGA samples as a surrogate marker of vtRNA expression (Corces et al., 2018). Although DNA methylation is a less direct and less accurate measurement of chromatin accessibility compared to ATAC-seq, 25 times more TCGA samples have been analyzed by DNA Methylation Arrays, including matched normal tissues samples, thus this dataset is of great significance for the current study. Our analyses revealed that vtRNA1-1 has accessible chromatin and small variation in all the evaluated tissues, which is consistent with its identity as the ancestor vtRNA gene and its significance as the major RNA component of the ubiquitous vault particle (Stadler et al., 2009; van Zon et al., 2001). In addition, vtRNA1-2, vtRNA1-3 and vtRNA2-1, have lower chromatin accessibility at their promoters and a larger variation among the samples. As expected, the average DNA methylation assessed by microarrays is negatively correlated with the chromatin accessibility assessed by ATAC-seq. Therefore, the landscape of vtRNA chromatin status was corroborated by both approaches. Yet, the low chromatin accessibility of vtRNA1-3 promoter is not reflected by a concomitant high DNA methylation, suggesting that additional regulatory features of this gene, acting in trans or cis, play a major influence in its chromatin structure. As was reported by Helbo et. al. using NOMe-seq assay (Nucleosome Positioning), the chromatin at vtRNA1-3 may be poised for transcription, and still require TFs to achieve physiological levels of transcription (Helbo et al., 2017).

The relative abundance of the 4 vtRNAs determined in the current study has been previously observed in various cell lines, including HSC CD34+, HL60 (Helbo et al., 2015), T24, colorectal RKO, human diploid fibroblasts IMR90 (Helbo et al., 2017) and MCF7 (Chen et al., 2018). Yet, others found different levels of the vtRNAs in MCF7, SW1573, GLC4, 8226, KB, SW620, MCF-7, HeLa, HEK-293 and L88/5 (van Zon et al., 2001), CBL, BL2 and BL41 (Nandy et al., 2009), EBV-negative Burkitt’s lymphoma cell lines and A549 (Li et al., 2015) and in LNCaP, PC3, DU145, RWPE-1, HEK, HeLa and MCF-7 (Stadler et al., 2009) cell lines, suggesting that the expression of vtRNAs may differ in laboratory cell lines compared to tissues. Alternatively, the methods used to quantify the vtRNAs are not comparable among the studies.

Seeking to investigate regulatory connections between the vtRNAs as a valuable tool to study their function, we determine the pairwise correlations between the promoters of the vtRNAs. The positive correlations found are compatible with a coordinated transcriptional control of the vtRNAs, that is greater for the three members of the vtRNA1 cluster (specially vtRNA1-1 and vtRNA1-3). The exception is the pair vtRNA1-2 and vtRNA2-1, whose chromatin accessibility and promoter methylation are not linked in the cohort and are indeed negatively associated in some comparisons.

As an alternative approach to investigate the vtRNA transcriptional co-regulation, we analyzed transcription and chromatin remodeling factors associated with their proximal promoter region. In agreement with their higher chromatin correlation, the vtRNA1 cluster have a larger core of common TFs (39) in comparison with vtRNA2-1 (23 TFs). The divergence of the transcriptional control of vtRNA2-1 was previously recognized given its unique regulatory elements and the complex developmental methylation it undergoes (Canella et al., 2010; Carpenter et al., 2018; Helbo et al., 2017; Nandy et al., 2009). Remarkably, the 23 TFs common to all vtRNAs are related to viral infections and cancer. The upregulation of the four vtRNAs upon viral infection has been reported for several viruses, so it is possible that this core of TFs participate in the coordinated expression of vtRNA during viral infection and related responses (Amort et al., 2015; Li et al., 2015; Nandy et al., 2009). These TFs regulate cell cycle and translational arrest as well as the inhibition of host innate immune response (Attar and Kurdistani, 2017; Ertosun et al., 2016; Horsley and Pavlath, 2002; Jean et al., 2000; Persengiev and Green, 2003; Rohini et al., 2018). Since these processes are also hallmarks of cancer, the dysregulation of the normal function of vtRNAs in viral response may provide a selective advantage for cancer cell to overcome the many sources of stress faced during neoplastic evolution. If that holds true, the core vtRNA TFs may be co-opted in the malignant context to tune vtRNA expression favoring tumor progression. In agreement with that hypothesis, ATF1 and ATF3 were associated with CREB response and increased cell viability (Persengiev and Green, 2003). Likewise, Golec et. al. 2019 showed that altered levels of vtRNA2-1 modulate PKR activation and consequently altered the levels of CREB phosphorylation during T cell activation, which is a prerequisite for IFN-g activity (Golec et al., 2019). Furthermore, E2F1, a cell cycle regulator, was described as a direct transcriptional regulator of vtRNA2-1 in cervical cancer cells (Li et al., 2017). Moreover, it was shown that MYC binding to vtRNA2-1 promoter raises its expression and could be the explanation to the increased levels of vtRNA2-1 in some tumors (Park et al., 2017). Similarly, TGFB1 provokes the demethylation of vtRNA2-1 promoter and consequently increases its expression in ovarian cancer (Ahn et al., 2018).

Due to its importance for cancer biology, vtRNA2-1 transcriptional regulation was recently more investigated (Yeganeh and Hernandez, 2020). Various studies reported tissue specific roles of vtRNA2-1 in normal (Golec et al., 2019; Lee et al., 2019a; Miñones-Moyano et al., 2013; Sallustio et al., 2016; Suojalehto et al., 2014) and cancer tissue (Ahn et al., 2018; Fort et al., 2018; Hu et al., 2017; Im et al., 2020; Jeon et al., 2012a; Kunkeaw et al., 2012; Lee et al., 2011; Lee, 2015; Lee et al., 2016, 2014a, 2014b; Lei et al., 2017; Li et al., 2017; Ma et al., 2020; Treppendahl et al., 2012) and epigenetic alterations of the locus have been described in different malignancies (Ahn et al., 2018; Cao et al., 2013; Fort et al., 2018; Helbo et al., 2015, 2017; Joo et al., 2018; Lee et al., 2014a, 2014b; Park et al., 2017; Romanelli et al., 2014; Treppendahl et al., 2012). Meanwhile, except for Kirsten Grønbæk group’s contributions on the chromatin characterization of vtRNA1-3 and vtRNA1-2 promoters (Helbo et al., 2015, 2017), little is known about the transcriptional regulation and expression of the other vtRNAs. Likewise, to our knowledge, the global landscape of vtRNA expression across different cancer types has not been investigated. Our study shows that, except for vtRNA1-1, which has a high promoter chromatin accessibility and low DNA methylation in all the tissues, the other vtRNA promoters exhibit different levels of chromatin accessibility and DNA methylation among the tissue types. The relative status of the chromatin in different tissues is accurately mirrored by both approaches in the tissues examined by ATAC-seq and Illumina DNA methylation arrays. Interestingly, the same analysis performed in the normal tissue samples reveals that the tissue specific variation in DNA methylation of vtRNA promoters in the neoplastic tissues is already established in their normal tissue counterparts, indicating that the regulation of vtRNA expression is tuned during normal development and cell differentiation.

While there is no apparent concerted modulation of the chromatin accessibility of the vtRNA promoters in most tissues, ACC and LGG tumors seem to be the exception. Concordantly, a global hypomethylation of malignant ACC tumors (Legendre et al., 2016; Rechache et al., 2012), and an aberrantly methylation processes associated to altered DNMTs activity in LGG tumors were reported (Nomura et al., 2019). However, more research is necessary to understand the molecular basis of these observations.

The differential expression of the vtRNAs from normal to primary tumor tissues revealed different patterns depending on tissue origin. The average DNA methylation of the vtRNA1-1 and vtRNA1-3 promoters is not significantly deregulated, whereas vtRNA1-2 and vtRNA2-1 are decreased and increased respectively in tumor tissues. The latter finding favors a candidate OG and TSG function of vtRNA1-2 and vtRNA2-1 in cancer respectively. Indeed, the epigenetic repression of vtRNA2-1 was previously reported by Romanelli et al. in BLCA, BRCA, COAD and LUSC analyzing the same data from TCGA (Romanelli et al., 2014). Furthermore, functional studies in LUSC, PRAD, ESCA and AML provided experimental support to that hypothesis (Cao et al., 2013; Fort et al., 2018; Lee et al., 2014a; Treppendahl et al., 2012). Yet, vtRNA2-1 has been also proposed as an OG in ovarian, thyroid, endometrial, cervical and renal cancer (Ahn et al., 2018; Hu et al., 2017; Lee et al., 2016; Lei et al., 2017; Li et al., 2017; Yeganeh and Hernandez, 2020). In agreement with the functional data found by Lei et. al. (Lei et al., 2017), we found an epigenetic de-repression of vtRNA2-1 promoter DNA methylation in renal carcinoma (KIRP). Likewise, a closer look at THCA samples, shows an average decrease in the DNA methylation of vtRNA2-1 promoter that supports the OG function described in this tissue (Lee et al., 2016). Unfortunately, normal ovarian, cervical and endometrial tissues are not available at the TCGA. Finally, vtRNA1-2 has an oncogenic pattern of inferred expression between normal and tumor tissues for 13 of 16 tissues analyzed, raising a possible oncogenic function for this RNA.

Our study found also an association of DNA methylation at the promoter with a shorter overall survival for vtRNA1-1, vtRNA1-2 and vtRNA2-1. This is surprising for vtRNA2-1, since it has both TSG and OG roles and TSG compatible expression profiles in 3 cancer types, inferred by DNA methylation of its promoter in tumor vs normal tissues and the average promoter DNA methylation is increased in tumor compared to normal tissue. A higher impact of vtRNA2-1 expression in patient survival in the oncogenic context may justify this discrepancy. On the contrary, the association between vtRNA1-2 low promoter DNA methylation and low patient survival is in agreement with its OG expression in cancer. The lack of survival association with the ATAC-seq values may be due to the small number of patients.

VtRNA expression in individual cancer types, inferred from the epigenetic status of their promoters, has been associated with patient survival in several studies of vtRNA2-1, one of vtRNA1-3 and none for vtRNA1-1 or vtRNA1-2. Low methylation or high expression of vtRNA2-1 promoter was associated with good prognosis or overall survival in lung (Cao et al., 2013), esophageal (Lee et al., 2014a), prostate (Fort et al., 2018), AML (Treppendahl et al., 2012), gastric (Lee et al., 2014b) and liver (Yu et al., 2020). Conversely, a worse prognosis or overall survival association of vtRNA2-1 was reported in thyroid (Lee et al., 2016) and ovarian (Ahn et al., 2018). From these reports, only prostate (Fort et al., 2018) and gastric (Lee et al., 2014b) studies used the TCGA data. Likewise, the methylation status of the vtRNA1-3 promoter associates with overall survival in the lower risk Myelodysplastic Syndrome patients (Helbo et al., 2015). Additionally, vtRNA1-1 and vtRNA1-2 correlate to chemotherapeutic resistance by direct interaction with drugs (doxorubicin, etoposide and mitoxantrone) and the modulation of vtRNA1-1 confirmed this finding in osteosarcoma cell lines (Gopinath et al., 2005, 2010; Mashima et al., 2008; van Zon et al., 2001). Furthermore, increased levels of vtRNA1-1 were associated with increased proliferation and chemoresistance due to GAGE6 induction in MCF-7 cells (Chen et al., 2018). Besides, Norbert Polacek group showed that vtRNA1-1 expression confers apoptosis resistance in several human cell lines (BL2, BL41, HS578T, HEK293, A549 and HeLa) and revealed it capacity to repressed intrinsic and extrinsic apoptosis pathway (Amort et al., 2015; Bracher et al., 2020). The later findings are in agreement with the lower patient survival associated with high levels of vtRNA1-1 expression cancer-wide observed in our analysis.

Seeking to get insights into the cancer related function of the vtRNAs we performed a genome wide search for genes co-regulated at the level of promoter chromatin accessibility. Remarkably, we identified immune and cytokine related terms for the genes co-regulated with vtRNA2-1. This association agrees with its proposed roles in innate immune modulation via PKR repression or OAS1 regulation and in cytokine production (Calderon and Conn, 2018; Golec et al., 2019; Lee et al., 2019b; Li et al., 2015). Furthermore, vtRNA2-1 has been associated with autoimmune disorders (Renauer et al., 2015; Weeding and Sawalha, 2018) and tumor engraftment in prostate cancer (Ma et al., 2020). This enrichment in viral infection pathways, together with the viral infection involvement of the core TFs common to all vtRNAs, reinforces the hypothesis of the re-utilization of a regulatory RNA induced upon viral response in favor of cancer development at upstream and downstream regulation steps. Additionally, we found a non-random clustering localization at chromosome 5 for genes co-regulated with vtRNA1-1. The chromosome arm 5q and regions chr5q31 and chr5q32-q33 were found frequently deleted in myelodysplastic syndrome (MDS) and acute myeloid leukemia (AML) (Fuchs, 2012; Treppendahl et al., 2012). Furthermore, the modulation of vtRNA2-1 and vtRNA1-3 was previously associated with human bone marrow CD34+ cells, AML and MDS (Helbo et al., 2015; Treppendahl et al., 2012). Yet, our analysis points to vtRNA2-1 and vtRNA1-2, and to a lesser extent to vtRNA1-1 and vtRNA1-3, as the most relevant players across cancer types.

Support for the involvement of the vtRNAs in the immune responses in the context of cancer came from the association of their chromatin status with the immune subtype categories defined by Thorsson et al. (Thorsson et al., 2018). Our findings suggest a possible role of the vtRNAs in lymphocyte response in tumor samples, since the immunologically quiet subtype shows the lowest leukocyte fraction and vtRNAs promoter chromatin accessibility (Thorsson et al., 2018). Meanwhile, the vtRNA2-1 and vtRNA1-2 expression distinguishes 2 groups among the remaining 4 subtypes. The participation of vtRNA2-1 in the induction of the IFN-g and IL-2 expression in activated T cells through PKR modulation (Golec et al., 2019) goes in agreement with this hypothesis. Although the same study demonstrated that vtRNA1-1 expression was unchanged during T cell activation, it is remarkable that the 5q31-q33 region, comprising the vtRNA1 locus and chromatin co-regulated genes, encodes several cytokines that regulate the differentiation of Th1 and Th2 lymphocytes and has been linked with susceptibility to infections (Jeronimo et al., 2007; Lacy et al., 2000; Naka et al., 2009; Rodrigues et al., 1999). Furthermore, vtRNA2-1 levels were recently associated with macrophages M1/M2 fates in prostate PC3 cell line mice xenografts via TGF-b (Ma et al., 2020), pathways that define the six immune subtypes (Thorsson et al., 2018). Finally, we found that the proliferation rate and wound healing of the tumors are associated with the levels of vtRNA1-2 chromatin accessibility, which reinforces a OG role of vtRNA1-2 in cancer. Nonetheless, the participation of the vtRNAs in immune cells response inside the neoplastic niche needs to be further investigated.

Taken together our analysis reveals the pattern of chromatin accessibility and DNA methylation at the four vtRNA promoters, analyzed by tissue of origin in the TCGA cohort. The agreement between both dataset and previous related literature endorse the use of these variables as surrogates of the vtRNA transcripts expression. The four vtRNAs seem to be co-regulated at the transcriptional level, and TFs involved in viral infection are likely to take part in their coordinated transcription. The pattern of expression in normal vs tumor tissue and the association with cancer cell pathways and patient survival suggest that vtRNA2-1 and vtRNA1-2 are possibly the more relevant contributors to cancer. The results favor tissue specific TSG and OG roles for vtRNA2-1, a still not investigated oncogenic role of vtRNA1-2 and a limited cancer driver effect of vtRNA1-1/3 in specific tissue types. Lastly, we uncovered new evidence linking the vtRNAs with the immune response, cell proliferation and overall survival in cancer, which guarantees further investigation.

## Materials and Methods

### ATAC-seq data

The ATAC-seq data obtained by The Cancer Genome Atlas (TCGA) consortium were retrieved from UCSC Xena Browser (Goldman et al., 2020) on date 03/10/2018. It comprises the genomic matrix TCGA_ATAC_peak_Log2Counts_dedup_promoter with normalized count values for 404 samples (385 primary solid tumor samples across 23 tissues). As is described in UCSC Xena Browser to calculate the average ATAC-seq values a prior count of 5 was added to the raw counts, then put into a “counts per million”, then log2 transformed, then quantile normalized; the result is the average value in the file (log2(count+5)-qn values) across all technical replicates and all biospecimens belonging to the same TCGA sample group. The gene promoter for ATAC-seq is defined as a region within −1000 to +100bp from the TSS site. Assignment of promoter peak to gene mapping information derives from the peak summit within the promoter region. Peak location information of the ATAC-seq values for 500 bp gene promoter regions was retrieved from the file TCGA_ATAC_Log2Counts_Matrix.180608.txt.gz. All the analyses were performed for tissues with at least five primary tumor samples (21 tissues), which resulted in the exclusion of CESC (2 primary tumor samples) and SKCM (4 primary tumor samples). Correlations between ATAC-seq and DNA promoter methylation were performed for the 329 primary tumors that are studied by both strategies (Supplementary Table 1).

### DNA methylation data

The DNA methylation data obtained from The Cancer Genome Atlas (TCGA) consortium was retrieved from UCSC Xena Browser (Goldman et al., 2020) on date 03/10/2018. It comprises the normalized beta-value of DNA methylation obtained using Illumina Infinium Human Methylation 450 BeadChip arrays of the total (9149 samples) normal adjacent tissues (746 samples across 23 tissues) and primary solid tumors (8403 samples across 32 tissues) of the Pan-Cancer TCGA cohort. The current study is limited to solid tumors since only AML DNA methylation is available at the TCGA. All the analysis was performed for tissues with at least five samples. Normal adjacent tissue comprises 16 tissues, excluding CESC (2 samples), GBM (2 samples), PCPG (3 samples), SARC (4 samples), SKCM (2 samples), STAD (2 samples) and THYM (2 samples). Primary tumors comprise 32 tissues. The normalized promoter average beta-values (500 bp bin) comprise the following CpG sites for vtRNA1-1: cg12532653, cg05913451, cg14633504, cg16615348, cg25602765, cg13323902, cg18296956; for vtRNA1-2: cg21161173, cg13303313, cg00500100, cg05174942, cg25984996, cg11807153, cg15697852; for vtRNA1-3: cg23910413, cg19065177, cg02053188, cg01063759, cg07379832, cg07741016; and for vtRNA2-1: cg16615357, cg08745965, cg00124993, cg26896946, cg25340688, cg26328633, cg06536614, cg18678645, cg04481923 (Supplementary Table 1).

### Immune Subtypes data

The data of Immune Subtypes defined by Thorsson et al. (Thorsson et al., 2018) is available at the Pan-Cancer TCGA and was retrieved from UCSC Xena Browser (Goldman et al., 2020) on date 03/10/2018 and from the supplementary material of Thorsson et al. article (Thorsson et al., 2018). ATAC-seq data of vtRNA promoters for the six Immune Subtype comprised a total of 364 samples: wound healing (105 samples), IFN-g dominant (93 samples), inflammatory (110 samples), lymphocyte depleted (43 samples), immunologically quiet (9 samples), TGF-b dominant (4 samples). DNA methylation average normalized beta-values of vtRNA promoters for the six Immune Subtype comprised 7693 samples including wound healing (1963 samples), IFN-g dominant (2137 samples), inflammatory (2126 samples), lymphocyte depleted (933 samples), immunologically quiet (383 samples), TGF-b dominant (151 samples) (Supplementary Table 5).

### Overall survival analysis

The Kaplan Meier curve analysis was performed with software GraphPad Prism 6 using Log-ranked Mantel-Cox test. We analyzed the ATAC-seq normalized values and DNA methylation average normalized beta-values for vtRNA promoters and survival data until 4000 days (99% of data). The log-rank statistics was calculated for quartiles of vtRNAs promoter chromatin accessibility comparison groups (25^th^ and 75^th^) (Supplementary Table 4).

### Pathway enrichment and cluster chromosome localization analyses

The genome wide correlation analysis of promoter chromatin accessibility (ATAC-seq normalized values) for each vtRNA in comparison with all the annotated genes was performed using the genomic matrix TCGA_ATAC_peak_Log2Counts_dedup_promoter retrieved from UCSC Xena Browser (385 primary tumor samples across 23 tissues) (Goldman et al., 2020) on date 03/10/2018. The genes showing a Spearman correlation r ≥ 0.4 were selected for pathway and cluster location analysis. The pathway enrichment analysis was done using STRING software for the Gene Ontology Biological Process and KEGG pathways categories (Szklarczyk et al., 2017). The cluster localization analysis was performed with two approaches, the Cluster Locator algorithm (Pazos Obregón et al., 2018) and the Enrichr software (Kuleshov et al., 2016).

### Statistical Analysis

Unless specified all the variables are expressed as average value ± standard deviation (SD). Statistical analyses were performed with one-way ANOVA for multiple comparison tests, including Sidak’s Honest Significant Difference test as a post-hoc test, as referred in the legend of the Figures. Spearman equations were applied to determine the correlations. All the analyses were done in R software (version 3.6) using libraries ggcorrplot, corrplot, xlsx, heatmap.2 (clustering distance measured by Euclidean and Ward clustering algorithms) and software GraphPad Prism 6. The statistical significance of the observed differences was expressed using the p-value (* p < 0.05, ** p < 0.01, *** p < 0.001, **** p-value < 0.0001). Differences with a p-value of < 0.05 were considered significant.

## Supplementary Materials: Supplementary Tables and Figure Legends

**Supplementary Figure 1.**
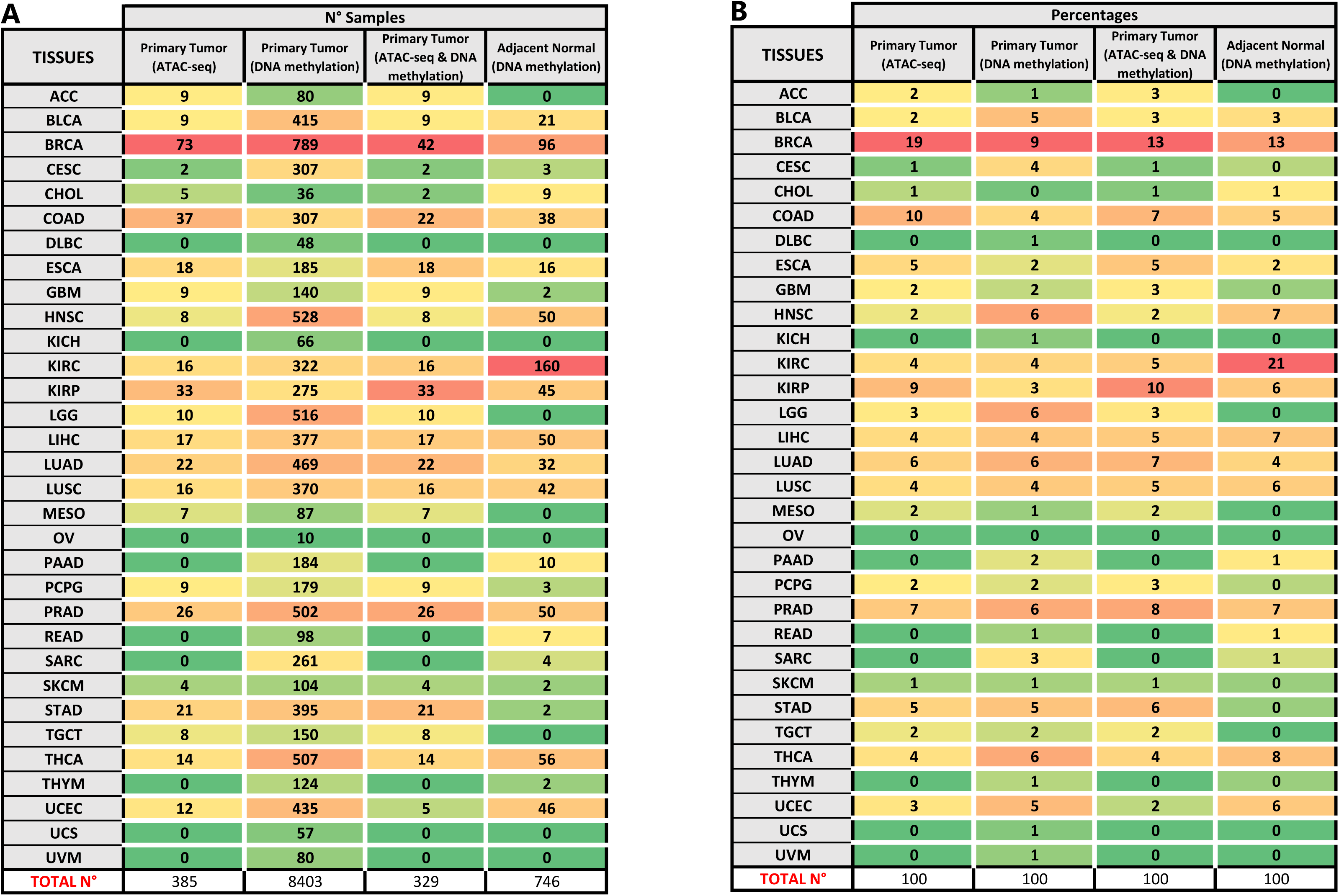
Data type and tissue sample distribution. Sample distribution per tissue type in the ATAC-seq, DNA methylation, ATAC-seq & DNA methylation and Normal adjacent tissues sample groups. **A.** Number of samples. **B.** Percentage of samples. The color reference, the number of samples of this tissue and the accounted percentage are presented to each tissue. Names references: ACC (Adrenocortical carcinoma), BLCA (Bladder Urothelial Carcinoma), BRCA (Breast invasive carcinoma), CESC (Cervical squamous cell carcinoma and endocervical adenocarcinoma), CHOL (Cholangiocarcinoma), COAD (Colon adenocarcinoma), DLBC (Lymphoid Neoplasm Diffuse Large B-cell Lymphoma), ESCA (Esophageal carcinoma), GBM (Glioblastoma multiforme), HNSC (Head and Neck squamous cell carcinoma), KICH (Kidney Chromophobe), KIRC (Kidney renal clear cell carcinoma), KIRP (Kidney renal papillary cell carcinoma), LGG (Brain Lower Grade Glioma), LIHC (Liver hepatocellular carcinoma), LUAD (Lung adenocarcinoma), LUSC (Lung squamous cell carcinoma), MESO (Mesothelioma), OV (Ovarian serous cystadenocarcinoma), PAAD (Pancreatic adenocarcinoma), PCPG (Pheochromocytoma and Paraganglioma), PRAD (Prostate adenocarcinoma), READ (Rectum adenocarcinoma), SARC (Sarcoma), SKCM (Skin Cutaneous Melanoma), STAD (Stomach adenocarcinoma), TGCT (Testicular Germ Cell Tumors), THCA (Thyroid carcinoma), THYM (Thymoma), UCEC (Uterine Corpus Endometrial Carcinoma), UCS (Uterine Carcinosarcoma) and UVM (Uveal Melanoma).

**Supplementary Figure 2.**
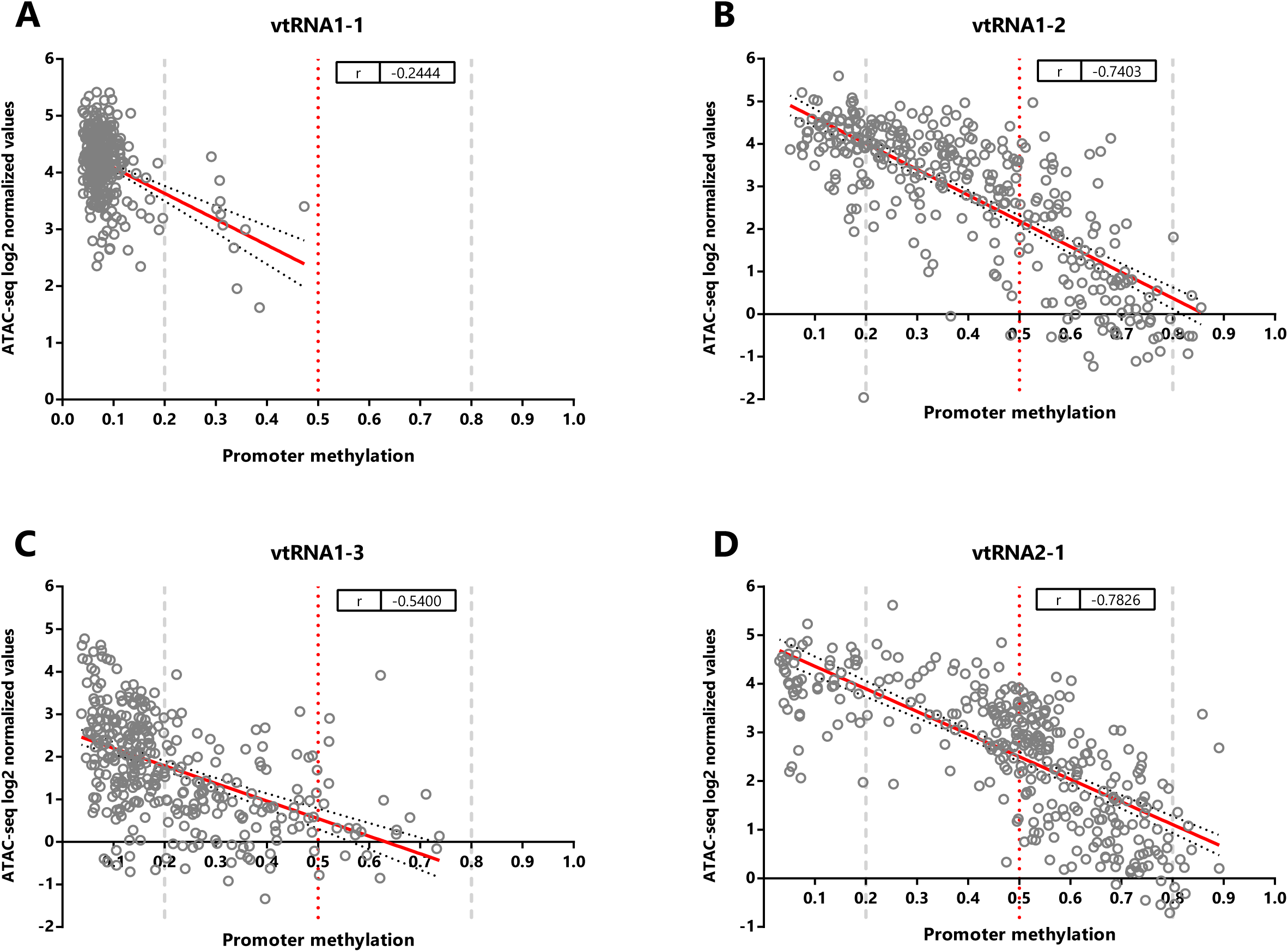
Correlation between ATAC-seq values and promoter methylation values of vtRNAs in Pan-Cancer TCGA dataset. **A-D.** The correlation was calculated between ATAC-seq data and DNA methylation average normalized beta values data for vtRNAs (vtRNAs1-1 (**A**), vtRNA1-2 (**B**), vtRNA1-3 (**C**) and vtRNA2-1 (**D**)), established in 329 primary tumors samples. The box plots show the median line and lower and upper quartile and the whiskers the 2.5 and 97.5 percentile. Horizontal grey striped and red dotted lines are shown to denote unmethylated (average beta-value ≤ 0.2), 50% methylated (average beta-value = 0.5) and highly methylated (average beta-value ≥ 0.8). Spearman correlation value and the best-fit line (red line) with 95% confidence bands (black dot lines) are shown.

**Supplementary Figure 3.**
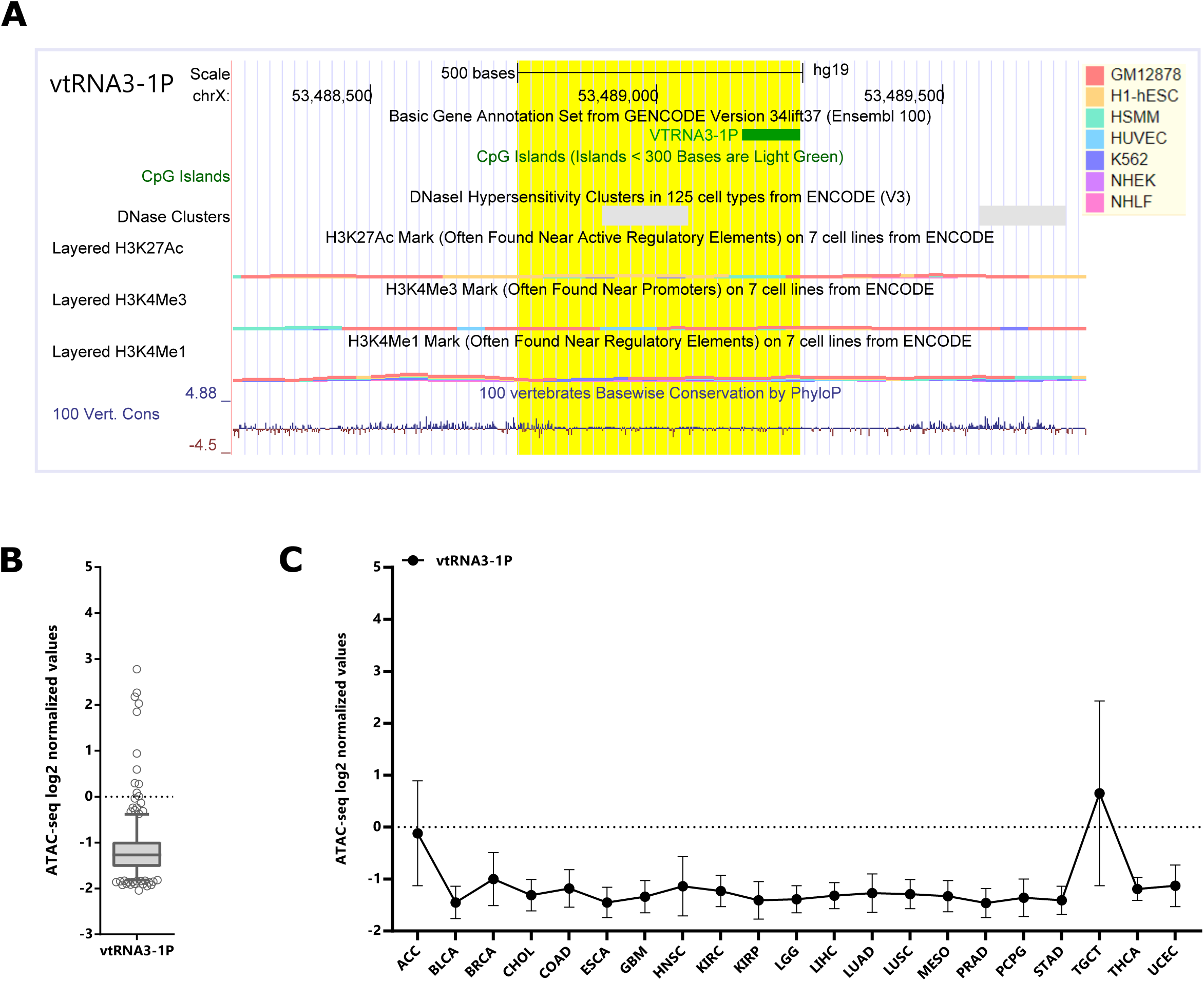
Genomic context of human vtRNA3-1P and the evaluated region of ATAC-seq. **A.** Genomic view of the 1.5 kb region of human vtRNA3-1P gene in UCSC Genome browser (GRCh37/hg19) centered at the 500bp of the ATAC-seq. Highlighted in *yellow* is the 500 bp ATAC-seq region used in the posterior analyses. The following Gene annotation and ENCODE Project tracks for seven cell lines (GM12878, H1-hESC, HSMM, HUVEC, K562, NHEK, NHL) are displayed: DNA accessibility (DNaseI hypersensitivity clusters (color intensity is proportional to the maximum signal strength)), DNA methylation (CpG islands length greater than 200 bp), histone modification (H3K27Ac, H3K4me1, H3K4me3 marks), conservation of the region in 100 Vertebrates (log-odds Phylop scores). The vertical viewing range of the tracks displays the default settings of the browser for each variable. **B.** The assay for transposase-accessible chromatin using sequencing (ATAC-seq) values for vtRNAs in different tumors present in PANCAN TCGA dataset (385 tumors samples across 23 cancer types) expressed as log2 normalized values. The box plots show the median and the lower and upper quartile, and the whiskers the 2.5 and 97.5 percentile of the distribution. **C.** The assay for transposase-accessible chromatin using sequencing (ATAC-seq) values for vtRNAs in 21 different tumors with at least 5 tumor samples present in Pan-Cancer TCGA dataset (385 tumors samples) expressed as log2 normalized values. The chart shows the average and standard error for each vtRNAs in each tumor.

**Supplementary Figure 4.**
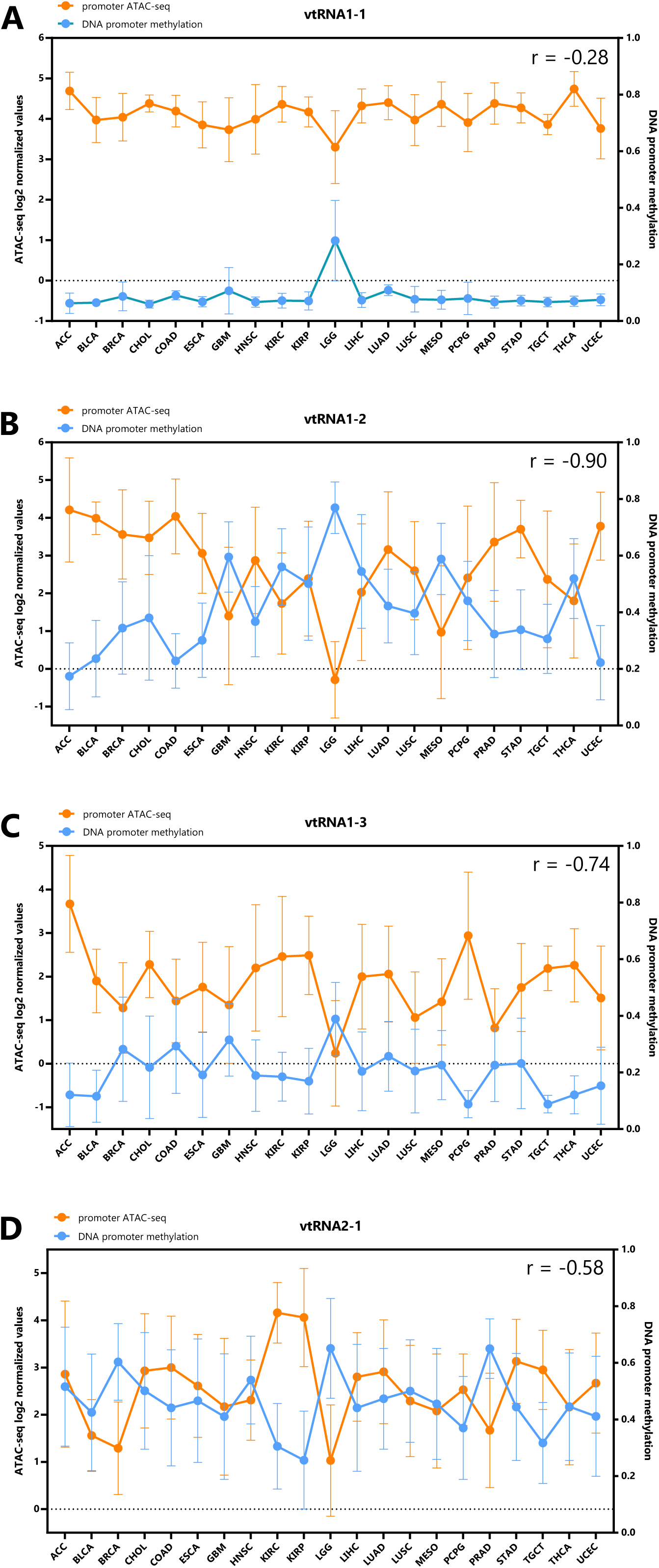
VtRNA promoter’s chromatin accessibility (ATAC-seq and DNA methylation) average values for tumor tissues from Pan-Cancer TCGA dataset. The average vtRNA promoters ATAC-seq and DNA methylation values for each primary tumor tissue type (21 tissues). VtRNA1-1 (**A**), vtRNA1-2 (**B**), vtRNA1-3 (**C**) and vtRNA2-1 (**D**). The Spearman correlation between averages ATAC-seq and DNA methylation values was calculated for each vtRNA. The chart shows the average and standard deviation for each tissue with at least five samples available at Pan-Cancer TCGA.

**Supplementary Figure 5.**
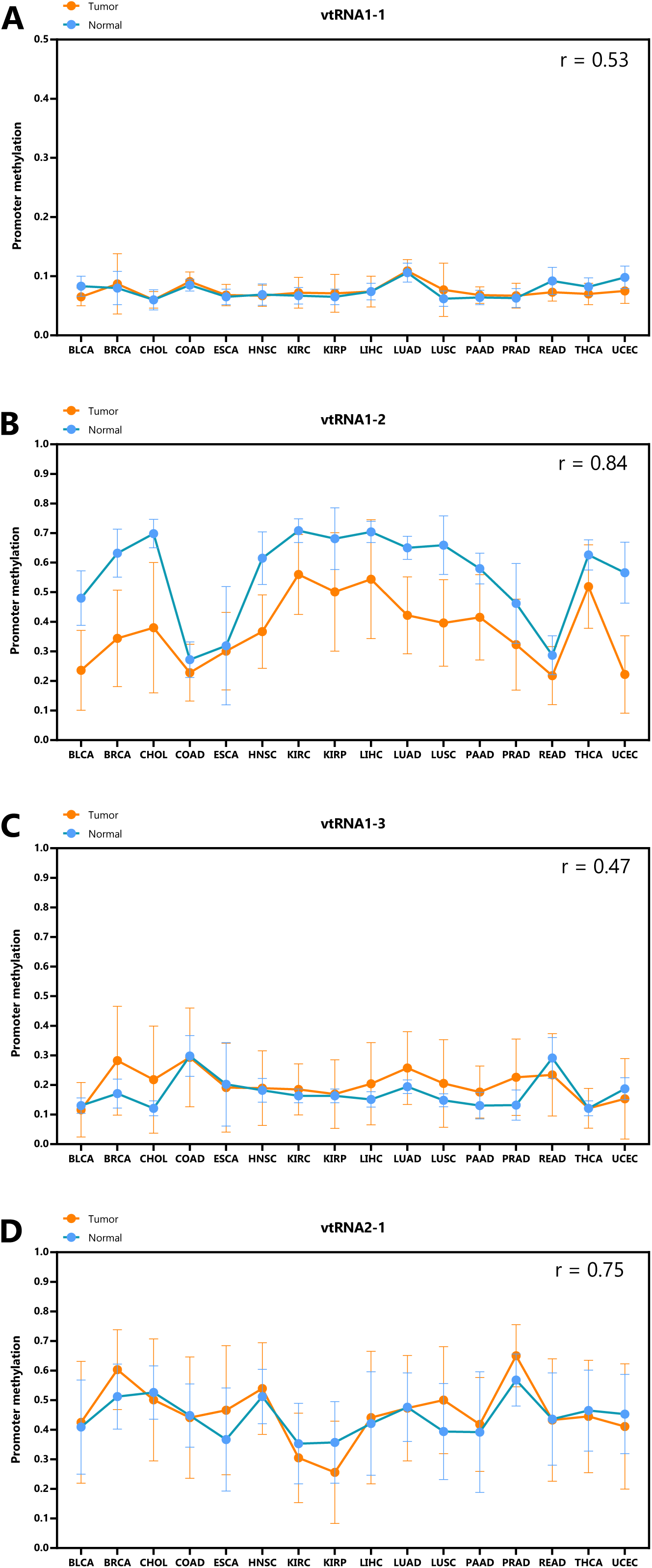
VtRNA promoter’s DNA methylation values for each normal and tumor tissues from Pan-Cancer TCGA dataset. The average promoter DNA methylation beta-values of normal adjacent (Normal) and all primary tumor (Tumor) tissues (16 tissue types) for vtRNA1-1 (**A**), vtRNA1-2 (**B**), vtRNA1-3 (**C**) and vtRNA2-1 (**D**) is shown. The Spearman correlation between tissue averages DNA methylation normalized beta values for each vtRNA was calculated. The chart shows the average and standard deviation for each tissue with at least five samples available at Pan-Cancer TCGA.

**Supplementary Table 1. ATAC-seq and DNA methylation data for vtRNA promoters in primary tumors and normal adjacent tissue samples.** Excel spreadsheets: ATAC-seq_data_500bp: All ATAC-seq data of vtRNAs promoter (500bp) data for primary tumor samples; DNA_methylation_500bp: All DNA methylation data and ATAC-seq data of vtRNAs promoter (500bp) data for total primary tumor and normal adjacent samples; DNA_methylation_NORMAL: All DNA methylation data of vtRNAs promoter (500bp) data for normal adjacent samples; DNA_methylation_TUMOR: All DNA methylation data of vtRNAs promoter (500bp) data for primary tumor samples; Normal_&_Tumor_matched: All DNA methylation data of vtRNAs promoter (500bp) data for primary tumor and normal adjacent samples.

**Supplementary Table 2. VtRNAs Transcription Factors Binding and KEEG enriched terms.** Excel spreadsheets: Binding_Factors: Transcription factors identified in the cell line K562 as ChIP-seq Peaks by ENCODE 3 project; KEEG_terms: enriched KEGG pathway terms (FDR < 0.05).

**Supplementary Table 3. DNA methylation, ATAC-seq data and associated survival data for primary tumors.** Excel spreadsheets: DNA-methylation_Survival_data: All DNA methylation data of vtRNAs promoter (500bp) and survival data for primary tumor samples; ATAC-seq_Survival_data: ATAC-seq data of vtRNAs promoter (500bp) and survival data for primary tumor samples.

**Supplementary Table 4. Correlation of ATAC-seq values between vtRNA and all genome promoters in primary tumor samples.** Excel spreadsheets: Spearman_correlation: Spearman correlation values of all promoter genes and vtRNAs in primary tumors samples; vtRNA1-1_cluster: Pathway enrichment and cluster chromosome localization data of vtRNA1-1; vtRNA1-2_cluster: Pathway enrichment and cluster chromosome localization data of vtRNA1-2; vtRNA1-3_cluster: Pathway enrichment and cluster chromosome localization data of vtRNA1-3; vtRNA2-1_cluster: Pathway enrichment and cluster chromosome localization data of vtRNA2-1.

**Supplementary Table 5. ATAC-seq and DNA methylation data for vtRNA promoters in primary tumors and the associated Immune Subtypes data.** Excel spreadsheets: DNA_methylation_data: All DNA methylation data of vtRNAs promoter (500bp) and Immune Subtypes data for primary tumor samples; DNA_methylation_Spearman_corr: Spearman correlation values of DNA methylation data of vtRNAs promoter (500bp) and Immune Subtypes data for primary tumor samples; ATAC-seq_data: All ATAC-seq data of vtRNAs promoter (500bp) and Immune Subtypes data for primary tumor samples; ATAC-seq_Spearman_corr: Spearman correlation values of ATAC-seq data of vtRNAs promoter (500bp) and Immune Subtypes data for primary tumor samples.

## Abbreviations

vtRNA: vault RNA
TCGA: The Cancer Genome Atlas
ATAC-seq: Assay for Transposase Accessible Chromatin with high-throughput sequencing\
NoMe-seq: Nucleosome Occupancy and Methylome sequencing
OG: Oncogene
TSG: Tumor Suppressor Gene
TCGA: The Cancer Genome Atlas
TSS: Transcription Start Site
Ave: Average
SD: Standard Deviation
ACC: Adrenocortical carcinoma
BLCA: Bladder Urothelial Carcinoma
BRCA: Breast invasive carcinoma
CESC: Cervical squamous cell carcinoma and endocervical adenocarcinoma
CHOL: Cholangiocarcinoma
COAD: Colon adenocarcinoma
DLBC: Lymphoid Neoplasm Diffuse Large B-cell Lymphoma
ESCA: Esophageal carcinoma
GBM: Glioblastoma multiforme
HNSC: Head and Neck squamous cell carcinoma
KICH: Kidney Chromophobe
KIRC: Kidney renal clear cell carcinoma
KIRP: Kidney renal papillary cell carcinoma
LGG: Brain Lower Grade Glioma
LIHC: Liver hepatocellular carcinoma
LUAD: Lung adenocarcinoma
LUSC: Lung squamous cell carcinoma
MESO: Mesothelioma
OV: Ovarian serous cystadenocarcinoma
PAAD: Pancreatic adenocarcinoma
PCPG: Pheochromocytoma and Paraganglioma
PRAD: Prostate adenocarcinoma
READ: Rectum adenocarcinoma
SARC: Sarcoma
SKCM: Skin Cutaneous Melanoma
STAD: Stomach adenocarcinoma
TGCT: Testicular Germ Cell Tumors
THCA: Thyroid carcinoma
THYM: Thymoma
UCEC: Uterine Corpus Endometrial Carcinoma
UCS: Uterine Carcinosarcoma
UVM: Uveal Melanoma

## Author Contributions

Conceptualization, M.A.D.; methodology, R.S.F. and M.A.D.; software, R.S.F.; formal analysis, R.S.F.; investigation, R.S.F. and M.A.D.; resources, M.A.D.; data curation, M.A.D and R.S.F.; writing—original draft preparation, R.S.F.; writing—review and editing, M.A.D.; visualization, R.S.F.; supervision, M.A.D.; project administration, M.A.D.; funding acquisition, M.A.D.

All authors have read and agreed to the published version of the manuscript.

## Funding

This study was funded by Agencia Nacional de Investigación e Innovación (ANII), Comisión Sectorial de Investigación Científica (CSIC), Comisión Académica de posgrado (CAP) and Programa de Desarrollo de las Ciencias Básicas (PEDECIBA) from Uruguay.

## Conflicts of Interest

The authors declare no conflict of interest.

## Notes

### Competing Interest Statement

The authors have declared no competing interest.

